# Microglia show differential transcriptomic response to Aβ peptide aggregates *ex vivo* and *in vivo*

**DOI:** 10.1101/2021.03.02.433544

**Authors:** Karen N. McFarland, Carolina Ceballos, Awilda Rosario, Thomas Ladd, Brenda Moore, Griffin Golde, Xue Wang, Mariet Allen, Nilüfer Ertekin-Taner, Cory C Funk, Max Robinson, Priyanka Baloni, Noa Rappaport, Paramita Chakrabarty, Todd E. Golde

## Abstract

Aggregation and accumulation of amyloid-β (Aβ) is a defining feature of Alzheimer’s disease (AD) pathology. To study microglial responses to Aβ, we applied exogenous Aβ peptide, in either oligomeric or fibrillar conformation, to primary mouse microglial cultures and evaluated system level transcriptional changes and then compared these to transcriptomic changes in the brains of CRND8 APP mice. We find that primary microglial cultures have rapid and massive transcriptional change to in response to Aβ. Transcriptomic responses to oligomeric or fibrillar Aβ in primary microglia, though partially overlapping, are distinct and are not recapitulated *in vivo* where Aβ progressively accumulates. Furthermore, though classic immune mediators show massive transcriptional changes in the primary microglial cultures, these changes are not observed in the mouse model. Together, these data extend previous studies which demonstrate that microglia responses *ex vivo* are poor proxies for *in vivo* responses. Finally, these data demonstrate the potential utility of using microglia as biosensors of different aggregate conformation, as the transcriptional responses to oligomeric and fibrillar Aβ can be distinguished.

## Introduction

Alzheimer’s disease (AD) is characterized by two hallmark pathologies, senile plaques containing amyloid-β (Aβ) aggregates and neurofibrillary tangles (NFTs) composed of hyperphosphorylated and aggregated tau. Amyloid plaques are the earliest manifestations of the disease process and can appear up to 20 years before the onset of cognitive symptoms (Bateman *et al*, 2012). Amyloid pathology, in the absence of tau or neurodegenerative pathology, defines pre-clinical AD and is the first step along the Alzheimer’s continuum in humans (Cummings, 2019; Jack *et al*, 2018; Vickers *et al*, 2016). In longitudinal studies, amyloid deposition precedes tau accumulation which is more closely tied to cognitive decline relative to amyloid (Hanseeuw *et al*, 2019; Villemagne *et al*, 2013). Furthermore, genetic data strongly support a causal, triggering role for aggregation and accumulation of Aβ in AD (Kunkle *et al*, 2019)— including the well-studied *APOE4* risk allele in late-onset AD which reduces the clearance of Aβ from the brain (Liu *et al*, 2013). Yet, despite intensive study, the precise mechanism by which accumulation of Aβ aggregates trigger the degenerative phase of the disease is not well understood.

As the primary immune and phagocytic cell in the brain, the role of microglia has been of growing interest in AD and other neurodegenerative disorders. “Resting” microglia, which constitute up to 10% of the brain, constantly sample the surrounding brain microenvironment and can rapidly respond to an insult (Aguzzi *et al*, 2013). In AD, the presence of increased “reactive” microglial cells both around senile plaques and in areas of neurodegeneration is a well-established pathological feature (Dickson, 1997; Dickson *et al*, 1988; Perlmutter *et al*, 1992). Notably, Aβ_42_ fibrils and oligomers cause microglia activation resulting in the release of pro-inflammatory cytokines which may contribute to neurotoxicity (Dewapriya *et al*, 2013; He *et al*, 2012; Jimenez *et al*, 2008; Wang *et al*, 2016). Alterations in microglial activation states can also impact both amyloid and tau pathology in varying ways that are dependent on both the stimulus, the model system and the pathology that is being assessed.

Over the last decade, a series of genetic studies has firmly linked microglial function to AD. Genetic studies of familial and late-onset AD implicate a large number of loci that contain immune genes in mediating risk for AD (Carrasquillo *et al*, 2017; Guerreiro *et al*, 2013; Harold *et al*, 2009; Jin *et al*, 2015; Jonsson *et al*, 2013; Kunkle *et al*., 2019; Lambert *et al*, 2009; Lambert *et al*, 2013; Sims *et al*, 2017). Furthermore, genetic studies identifying coding variants in three microglial-specific genes (PLCG2, ABI3 and TREM2) highlight the important role microglia play during neurodegeneration (Bellenguez *et al*, 2017; Conway *et al*, 2018; Guerreiro *et al*., 2013; Jin *et al*, 2014; Jonsson *et al*., 2013; Sims *et al*., 2017; Strickland *et al*, 2020; van der Lee *et al*, 2019). Additionally, systems level data analysis of spatial, single-cell, single-nuclei as well as bulk RNA-sequencing (RNA-seq) studies reveal perturbations in immune transcriptional networks as well as distinct subpopulations of microglia that are perturbed in the AD brain (Chen *et al*, 2020; Conway *et al*., 2018; Friedman *et al*, 2018; Hammond *et al*, 2019; Keren-Shaul *et al*, 2017; Krasemann *et al*, 2017; Li *et al*, 2019; Olah *et al*, 2020).

The study of microglial cells is challenging in that they are highly responsive to external stimuli and rapidly alter their phenotype once removed from the brain (Bennett *et al*, 2016). Indeed, systems level transcriptomic studies show that primary microglial cells are poor proxies for *in vivo* microglia (Butovsky *et al*, 2014). Even rapid isolation of microglial and subsequent “omic” analyses can be challenging as it is clear the isolation process is sufficient to induce some transcriptional—and likely functional changes. Nevertheless, many labs—including our own—study primary microglial cells in culture. In particular, the application of exogenous Aβ aggregates to microglial is a widely used methodology to study both how microglial respond to Aβ and how effectively the microglia can phagocytose and degrade Aβ.

Here we used RNA-seq to examine the systems level response of primary microglia in culture to synthetic Aβ_42_ aggregates in either oligomeric (oAβ) or fibrillar (fAβ) form. Our analyses of the transcriptomic data show that microglial cells in culture show massive transcriptional changes when challenged with Aβ_42_ aggregates. Though some of the differentially expressed genes in response to the different forms of Aβ_42_ are altered similarly, many show differential expression in response to oAβ or fAβ. We also compared this global transcriptional response to Aβ_42_ in primary microglial cells in culture to transcriptomic data from a mouse model of amyloid deposition—the APP transgenic CRND8 mouse— at 3 to 20 months of age (Chishti *et al*, 2001). Subsequent comparisons of these datasets indicate that most Aβ transcriptional responses in microglia are largely not replicated in the intact brain. This comparison demonstrates that the transcriptional response to Aβ in primary cultures poorly reflect the response to Aβ by microglial cells in the mouse brain. These data amplify the message of several other recent studies indicating that one must be very cautious when using primary microglial cells cultured in isolation to infer mechanistic insights about microglial function *in vivo*.

## Results

### Large Transcriptomic Changes in primary microglia following Aβ treatment

Pre-formed oligomeric (oAβ) or fibrillar (fAβ) forms of Aβ_42_ peptide were applied to primary microglia cultures for 1- or 12-hr (Figure 1A). oAβ and fAβ were characterized by Western blot and a representative image is shown (Figure 1B) demonstrating differences in the high molecular weight species between oligomeric and fibrillar Aβ preparations. Following treatment, RNA was isolated and sequenced to identify transcriptional changes in primary microglia that are responsive to different conformations of Aβ_42_ peptide. As noted in the methods, this data along with the mouse CRND8 RNAseq data is publicly available and can be viewed using an interactive data portal. Using cut-off values of a p-value (adjusted for multiple comparisons) ≤ 0.05 and an absolute log2 fold-change of 0.5, we identified acute transcriptional changes following fAβ application after just 1-hr (versus control) with 997 upregulated and 960 downregulated genes (Figure 2A, Supplemental Data 1). Gene ontology (GO-MF) and KEGG pathway analysis identified downregulated genes as being enriched in cytoskeletal and extracellular matrix organization (i.e., *tubulin binding, motor activity*, *extracellular matrix structural constituent*; Figure 2E, Supplemental Data 1) while upregulated genes were involved in immune system responses (i.e., *RAGE receptor binding, chemokine activity)* and kinase activity (i.e., *MAP kinase phosphatase activity*; Figure 2F, Supplemental Data 1).

**Figure 1.**
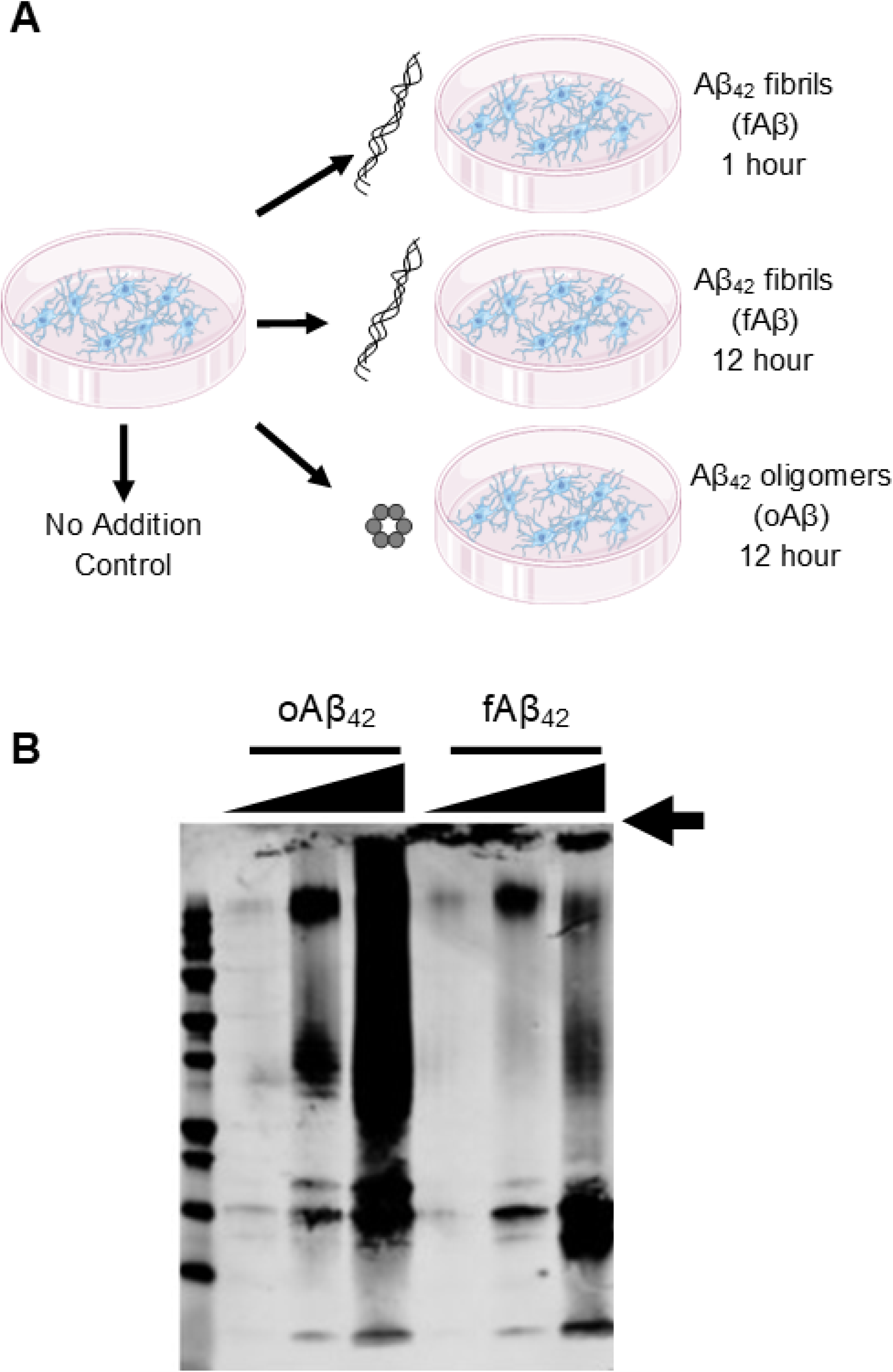
Aβ treatments of primary microglial cultures. A) Experimental paradigm for the applications of Aβ conformational species onto primary microglia cell cultures. B) Western blot analysis with anti-Aβ antibody (6E10) of representative oAβ and fAβ preparations. The oAβ preparation shows a smear with characteristic banding patterns while the fAβ preparation contains a significant amount of Aβ that do not enter the gel (arrow).

**Figure 2.**
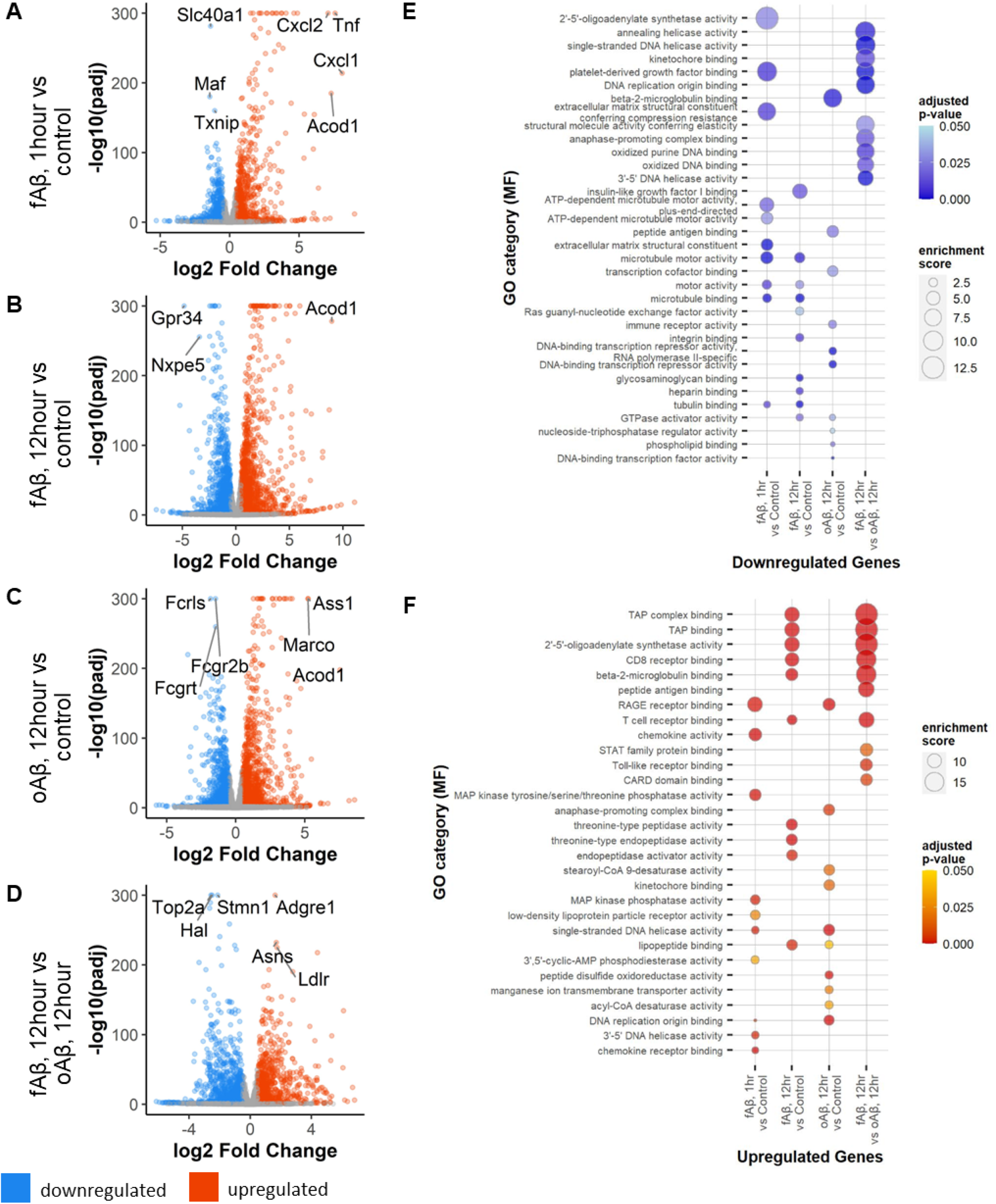
Differential gene expression in primary microglia following treatment with Aβ42 oligomers (oAβ) or fibrils (fAβ). A) Total changes in down- (blue) and up- (red) regulated genes in primary microglia following 1-hr of fAβ treatment versus control. B) Volcano plot of DEGs after 12-hr of fAβ42 treatment versus control in primary microglial cultures. C) Volcano plot of DEGs after 12-hr of oAβ42 treatment in primary microglial cultures. D) Volcano plot of DEGs after 12-hr of Aβ42 fibrils treatment in primary microglial cultures. E) Bubble plots of GO category enrichment results for downregulated genes. F) Bubble plots of GO category enrichment results for upregulated genes. Plots for GO category over-enrichment analyses show the top 10 hits for each comparison by enrichment score following a filter step by a p-value adjusted for multiple comparisons of ≤ 0.05 and keeping GO categories with greater than 5 genes within the category.

After 12-hr of fAβ treatment, we identified 1,755 upregulated and 1,975 downregulated genes when compared with control (Figure 2B, Supplemental Data 2). GO and KEGG pathway analysis revealed enrichment of downregulated genes involved in cytoskeletal and extracellular matrix organization (i.e., *tubulin binding, motor activity*, *extracellular matrix structural constituent)* in addition to *heparin binding* and *glycosaminoglycan binding* (Figure 2E, Supplemental Data 2). Genes upregulated after 12-hr fAβ treatment were enriched in genes involved with antigen processing (TAP binding) and proteolytic activity (i.e., *endopeptidase activator activity, threonine-type peptidase activity;* Figure 2F, Supplemental Data 2).

We next examined the effect of a 12-hr oAβ treatment (versus control) on primary microglial cultures. We identified 1,608 upregulated and 1,394 downregulated genes after 12-hr of oAβ (Figure 2C, Supplemental Data 3). GO and KEGG pathway analysis revealed that downregulated genes are primarily involved in DNA transcription (i.e., *DNA-binding transcriptional repressor activity, transcription cofactor binding;* Figure 2C, 2E, Supplemental Data 3). Genes upregulated by oAβ treatment are enriched with GO terms suggestive of cell cycle involvement (i.e., *anaphase-promoting complex binding, kinetochore binding*; Figure 2F, Supplemental Data 3). A number of the top GO category hits overlap somewhat between the 1- and 12-hr fAβ treatments; however, many of the changes seen following oAβ treatment stand in stark contrast to those seen following both fAβ treatments.

To further examine differences and similarities in transcriptional changes between fAβ and oAβ treatments, we directly compared gene expression at 12-hr of fAβ treatment (numerator) against gene expression at 12-hr of oAβ treatment (denominator) to identify differentially expressed genes in these conditions. This comparison revealed disparate changes in transcriptional responses between the conformations of Aβ peptide and identified 982 upregulated genes and 1,348 downregulated genes in fAβ versus oAβ treatments (Figure 2D, Supplemental Data 4). Affected downregulated genes (down in fAβ while up in oAβ) primarily affected cell cycle and DNA binding activities (i.e., *DNA replication origin binding, kinetochore binding;* Figure 2E, Supplemental Data 4) while upregulated genes (up in fAβ relative to oAβ) were enriched in immune system responses (i.e., *TAP binding, T cell receptor binding*; Figure 2F, Supplemental Data 4).

### Primary Microglia have unique transcriptional responses to Aβ conformations

To directly identify disparate changes in transcription in response to Aβ conformation, we compared the log-fold changes for differentially expressed genes in these Aβ treatments (Supplemental Figure 1). We find a strong correlation (R = 0.74) when comparing treatments of fAβ, 12-hr (versus control) against oAβ, 12-hr (versus control). The 865 commonly upregulated genes are enriched with GO terms involved with peptidase and chemokine activity (i.e., *threonine-type endopeptidase activity*) while the 865 commonly downregulated genes are enriched in terms involving post-translational modifications (*histone demethylase activity, ubiquitin-like protein ligase activity*) (Supplemental Figure 1B, Supplemental Data 5). Interestingly, the 170 genes that are upregulated in oAβ, 12-hr treatment but downregulated in fAβ, 12-hr treatments are involved in cell cycle (i.e., *anaphase-promoting complex*) and microtubule motor activities (i.e., *motor activity, ATP-dependent microtubule motor activity*). The 51 genes downregulated in oAβ, 12-hr treatment but upregulated in fAβ, 12-hr treatment which are involved in antigen binding and immune responses (i.e., *TAP complex binding, CD8 receptor binding*).

An analysis comparing fAβ, 1-hr treatment with oAβ, 12-hr treatment reveals similar results (Supplemental Figure 1C, D, Supplemental Data 6). Commonly upregulated genes (515 genes) have roles involving the immune system (*RAGE receptor binding*, *chemokine activity*) and kinase activities (*MAP kinase tyrosine/threonine phosphatase activity*) while there was no significant enrichment of GO terms (p-value adjusted for multiple comparisons ≤ 0.1) for the 387 commonly downregulated genes. The divergently responding 56 genes that are upregulated in oAβ, 12-hr treatment but downregulated in fAβ, 1-hr treatment are involved in the cell cycle (*anaphase-promoting complex binding*) and microtubule motor processes (*microtubule motor activity*) while the 78 genes upregulated in fAβ, 1-hr treatment, but downregulated in fAβ, 12-hr treatment are involved in the innate immune response (*complement component C1q complex binding*). This analysis highlights the upregulation of genes involved in the cell cycle and microtubule motor pathways following oAβ treatment.

A direct comparison of significant changes in gene expression between acute 1-hr versus longer-term 12-hr fAβ treatments expose 515 commonly upregulated genes involved in cytokine and immune activity (*immunoglobulin receptor binding, chemokine receptor binding)* and 507 commonly downregulated genes involved in microtubule motor activity (*microtubule binding, microtubule motor activity*; Supplemental Figure 1E, F, Supplemental Data 7). Longer-term fAβ treatment resulted in an upregulation of 93 genes that are initially downregulated in 1-hr fAβ treatment that are involved in adenylation and GTPase activities (*adenylyltransferase activity, GTPase activity, nucleoside-triphosphatase activity*). Acute fAβ, 1-hr treatment triggered an upregulation of 98 genes that are downregulated after 12-hr of treatment which are enriched in diverse GO terms including *complement component C1q complex binding, DNA helicase activity*, and *integrin binding*.

For a more comprehensive view of these disparate changes in microglia following the application of different Aβ species, we examine all genes that were identified as a differentially expressed gene (DEG) in any of the of the three treatment paradigms versus control and plotted a heatmap of their z-scores with hierarchical clustering of the genes (Figure 3). Clear patterns of transcriptional changes can be seen between conditions. To identify the genes within these clusters, we cut the hierarchical tree at a height of 5.75 which resulted in 13 gene clusters that were then analyzed by GO analysis (Figure 3 and Table 1, Supplemental Data 8). By this analysis, we identified clusters of genes that have similarities in expression patterns following treatment with different Aβ conformations. For example, genes in cluster 10 are involved in transcriptional processes and have decreased expression in all three conditions compared with controls. However, this analysis also highlights the clusters of genes that have a unique transcriptional signature in response to specific Aβ conformations. Genes within clusters 11 & 9 have increased expression levels following acute fAβ treatment and are enriched in terms involving metabolic processes as well as immune responses and cell signaling. Genes in cluster 7 are increased following long-term fAβ treatment and encompass functions of the antigen processing and the immune system. Genes in clusters 1 & 5 are strongly increased in expression after oAβ treatment and are involved in cell cycle and nucleobase metabolism.

**Figure 3:**
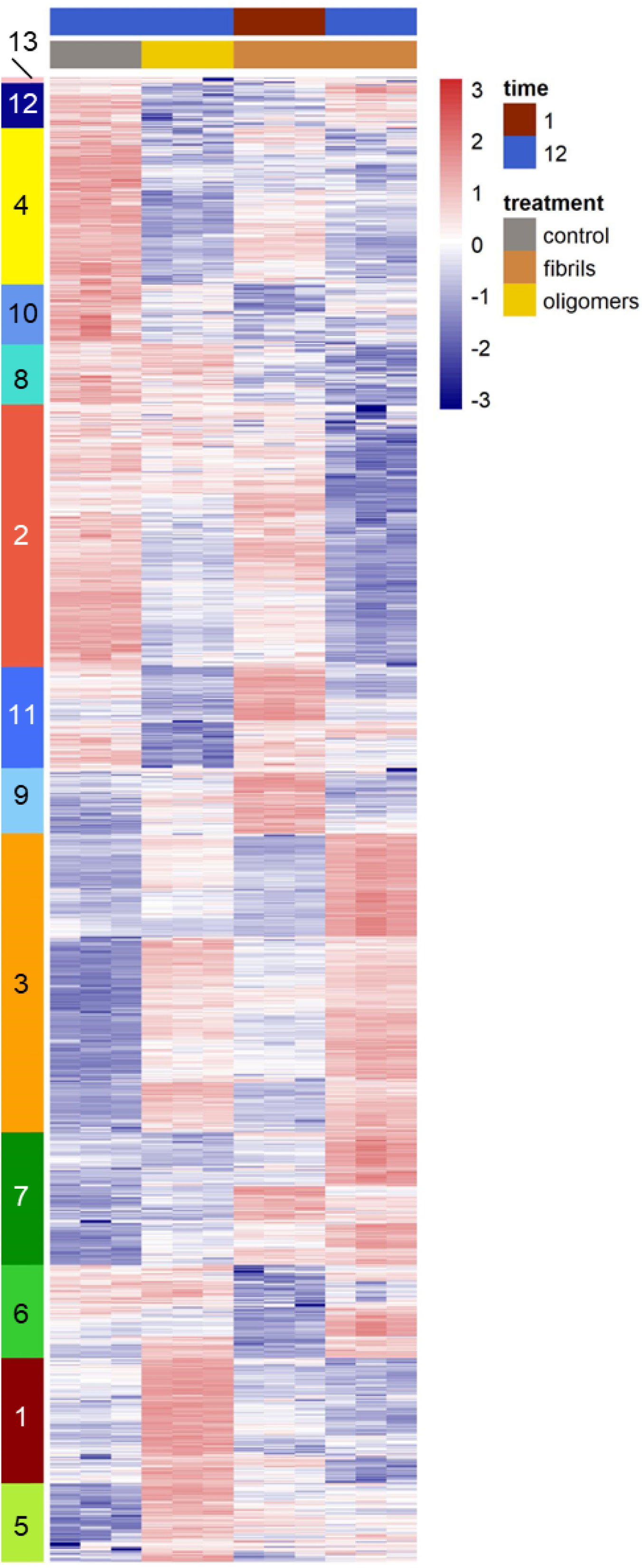
Hierarchical clustering of differentially expressed genes in Aβ42-treated microglia reveal unique gene signatures. Hierarchical clustering Z-scores of gene expression data. A cut height of h = 5.75 was applied to identify clusters of genes with similar expression patterns which produced 13 clusters.

**Table 1:** GO analysis of differentially expressed gene clusters in Aβ-treated microglia. GOseq analysis to analyze GO category over-enrichment was applied to these clusters identified in Figure 3. The top 3 categories are shown after ranking by enrichment score and filtering for genes with a p-value of less than 0.05 and to remove categories with less than 5 genes within the category.

| Cluster | Control | oAβ, 12-hr | fAβ, 1-hr | fAβ, 12-hr | GO category (MF) | Enrichment score | P-value |
| --- | --- | --- | --- | --- | --- | --- | --- |
| 13 | ↑ | mix | mix | mix | carbohydrate:proton symporter activity | 24.95 | 9.15E-03 |
|  |  |  |  |  | JUN kinase kinase kinase activity | 22.46 | 1.02E-02 |
|  |  |  |  |  | neurexin family protein binding | 17.28 | 1.32E-02 |
| 12 | ↑ | ↓ | ↓ | ↑ | kainate selective glutamate receptor activity | 5.44 | 4.14E-02 |
|  |  |  |  |  | testosterone dehydrogenase (NAD+) activity | 5.44 | 4.24E-02 |
|  |  |  |  |  | alpha-adrenergic receptor activity | 5.44 | 4.26E-02 |
| 4 | ↑ | ↓ | ↑ | ↓ | ATPase inhibitor activity | 4.66 | 3.16E-04 |
|  |  |  |  |  | RNA polymerase II transcription cofactor binding | 3.11 | 9.28E-03 |
|  |  |  |  |  | LBD domain binding | 2.66 | 1.26E-02 |
| 10 | ↑ | ↓ | ↓ | ↓ | histone demethylase activity (H3-K9 specific) | 8.02 | 3.99E-06 |
|  |  |  |  |  | leucine binding | 8.02 | 1.64E-03 |
|  |  |  |  |  | insulin binding | 8.02 | 2.14E-03 |
| 8 | ↑ | ↑ | ↓ | ↓ | DNA insertion or deletion binding | 8.09 | 1.49E-03 |
|  |  |  |  |  | MutLalpha complex binding | 8.09 | 1.52E-03 |
|  |  |  |  |  | sphingosine N-acyltransferase activity | 6.94 | 2.06E-03 |
| 2 | ↑ | ↓ | ↑ | ↓ | 3-hydroxyacyl-CoA dehydrogenase activity | 2.08 | 4.12E-03 |
|  |  |  |  |  | co-receptor binding | 1.85 | 1.52E-03 |
|  |  |  |  |  | dolichyl-phosphate-mannose-protein mannosyltransferase activity | 1.85 | 6.07E-03 |
| 11 | ~ | ↓ | ↑ | ↓ | STAT family protein binding | 1.39 | 7.17E-03 |
|  |  |  |  |  | complement component C1q binding | 1.39 | 7.36E-03 |
|  |  |  |  |  | TAP complex binding | 1.24 | 9.57E-03 |
| 9 | ↓ | ~ | ↑ | ↓ | stearoyl-CoA 9-desaturase activity | 7.39 | 1.72E-03 |
|  |  |  |  |  | MAP kinase tyrosine/serine/threonine phosphatase activity | 6.83 | 8.84E-06 |
|  |  |  |  |  | acyl-CoA desaturase activity | 6.34 | 2.39E-03 |
| 3 | ↓ | ↑ | ↓ | ↑ | threonine-type endopeptidase activity | 3.25 | 4.46E-14 |
|  |  |  |  |  | threonine-type peptidase activity | 3.25 | 4.46E-14 |
|  |  |  |  |  | proteasome-activating ATPase activity | 3.25 | 7.92E-05 |
| 7 | ↓ | ↓ | mix | ↑ | TAP binding | 6.99 | 7.76E-10 |
|  |  |  |  |  | TAP complex binding | 6.10 | 6.10E-07 |
|  |  |  |  |  | CD8 receptor binding | 5.99 | 4.93E-08 |
| 6 | mix | mix | ↓ | ↑ | platelet-derived growth factor binding | 6.51 | 6.36E-07 |
|  |  |  |  |  | extracellular matrix structural constituent conferring compression resistance | 5.58 | 1.57E-06 |
|  |  |  |  |  | cobalt ion binding | 4.69 | 4.11E-04 |
| 1 | ~ | ↑ | ↓ | ↓ | single-stranded DNA-dependent ATPase activity | 7.97 | 3.77E-18 |
|  |  |  |  |  | kinetochore binding | 7.76 | 3.05E-06 |
|  |  |  |  |  | single-stranded DNA-dependent ATP-dependent DNA helicase activity | 7.28 | 2.24E-07 |
| 5 | ↓ | ↑ | ~ | ~ | G protein-coupled adenosine receptor activity | 5.31 | 3.35E-03 |
|  |  |  |  |  | structural constituent of presynapse | 4.13 | 5.78E-03 |
|  |  |  |  |  | low-density lipoprotein particle receptor activity | 3.71 | 6.96E-03 |

### Gene network changes in microglia highlight specific transcriptional responses to Aβ conformations

We applied a weighted gene co-expression network analysis (WGCNA) onto the expression data from Aβ-treated primary microglial cultures. WGCNA is a method to study biological networks by analyzing pair-wise correlations between the genes within the dataset (Langfelder & Horvath, 2008). We identified seventy-one co-expression modules (Figure 4, Supplemental Data 9). We correlated the modules with treatment paradigms (Figure 4A) and annotated these modules using a gene overlap analysis (Shen, 2020) with genes identified with sub-populations of microglial cells identified in prior bulk, single-cell (sc-), single-nuclear (sn-) RNA-seq or spatial transcriptomic studies (Figure 4B; Supplemental Data 10). We additionally annotated the modules by KEGG and GO analysis to identify enrichment of pathways within the modules (Figure 4C, Supplemental Data 11). By relating the modules to each treatment condition, we observed interesting patterns in module behavior.

**Figure 4:**
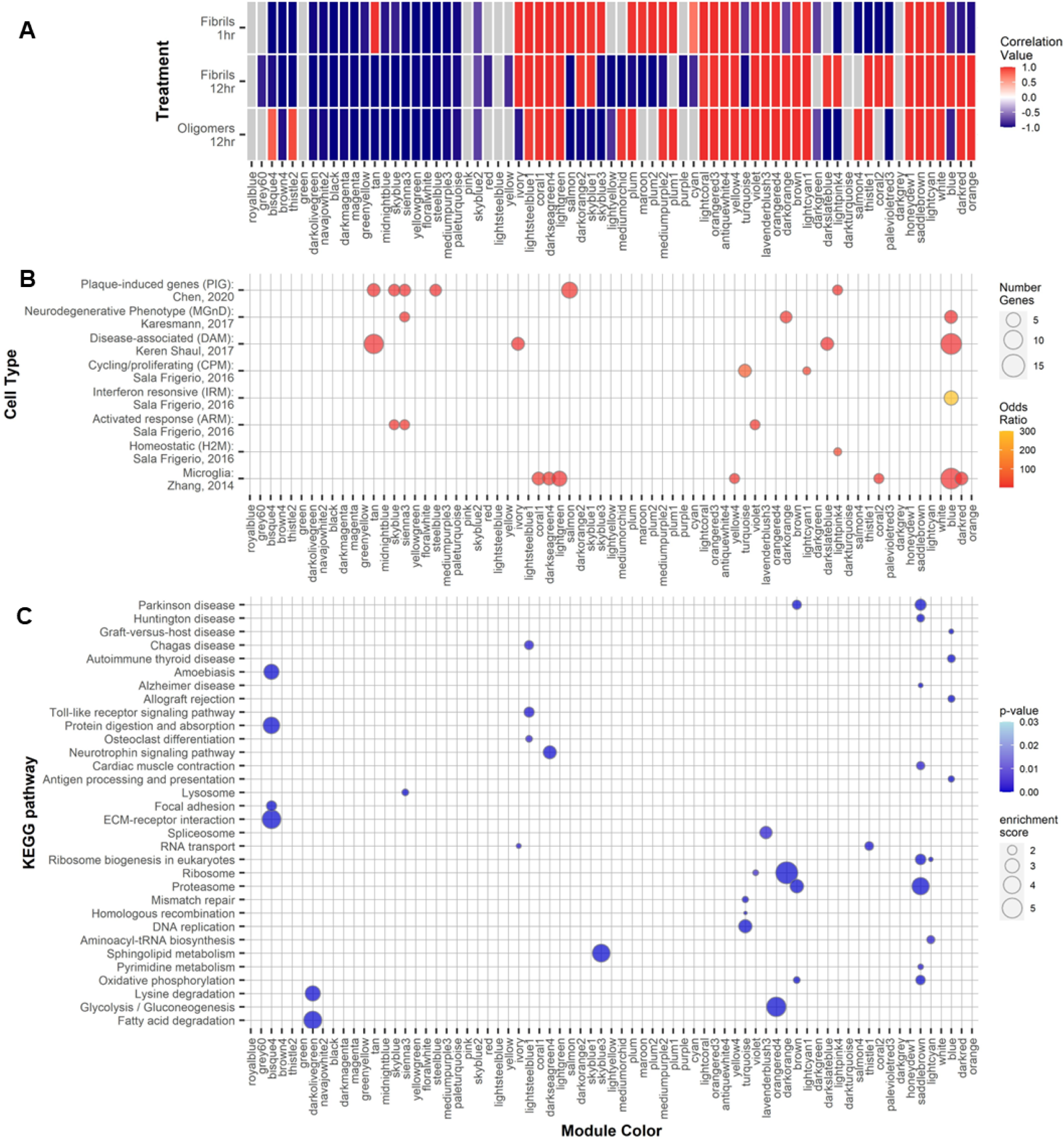
Weighted Gene Correlation Network Analysis (WGCNA). Gene modules found by WGCNA in Aβ-treated primary microglia. A) Modules are colored in a heatmap by their correlation value with the different Aβ treatments. Modules with non-significant p-values associated or with an absolute correlation value or less than 0.5 are indicated in grey. B) Bubble plot of a gene overlap analysis to identify shared genes between the module and previously identified microglial sub-types. Modules with significant (p ≤ 0.05) odds-ratios of overlapping genes are colored as in the scale to the right. The number of overlapping genes is indicated by the dot size. C) KEGG pathway over-enrichment analysis for genes within each module. Pathways with an over-represent p-value ≤ 0.05, the number of module genes within the pathway > 5 and an enrichment score > 1.5 are depicted. P-value is indicated by the color scare and the enrichment score by the dot size.

Of these modules, seventeen modules are positively correlated with all forms of treatment and indicate a non-specific response to Aβ treatment. These modules include antiquewhite4, brown, coral1, darkseagreen4, honeydew1, lavenderblush3, lightcoral, lightcyan, lightcyan1, lightgreen, lightsteelblue1, orangered3, orangered4, saddlebrown, violet, white and yellow4. GO and KEGG pathway analysis reveals that genes within these modules are involved in a variety of molecular functions previously linked with AD including cytokine and chemokine activities (lightgreen) the proteosome (brown and saddlebrown), the splicesome (lavenderblush3 and saddlebrown) as well as neurodegenerative pathways including AD, Parkinson’s disease, Huntington’s disease (brown and saddlebrown). Fifteen modules are negatively correlated with all forms of Aβ treatment—again, indicating a non-specific response—and include the black, brown4, darkmagenta, darkolivegreen, floralwhite, greenyellow, magenta, mediumpurple3, midnightblue, navajowhite2, paleturquoise, sienna3, skyblue, skyblue2, steelblue and yellowgreen modules. The genes within these modules are enriched in genes involved in Rab and Ras GTPase activities (mediumpurple3) and with fatty acid metabolism (darkolivegreen).

Six modules are positively correlated with acute, 1-hr fAβ treatment and are either negatively correlated or not significantly correlated with the other treatments. These modules characterize the acute response to fAβ treatment and include the tan, salmon, skyblue3, maroon, plum2 and cyan modules. These modules represent genes with functions involved with ion channel activities (cyan), histone modification activity (plum2), RNA processing and splicing, protein ubiquitination and acetylation (skyblue3).

There are five modules that are positively correlated to long-term, 12-hr fAβ and include the darkslateblue, lightpink4, palevioletred3, blue and coral2 modules. Genes within these modules are enriched with genes with immune/inflammatory/cytokine functions (blue), RNA binding (darkslateblue), GTPase activity (palevioletred and coral2) and transcriptional regulation (lightpink4). Additionally, it is within the blue module that the majority of reactive and responsive microglial markers reside (Figure 4C).

Five other modules are positively correlated with long-term, 12-hr oAβ treatment and include bisque4, thistle2, mediumorchid, turquoise and salmon4. These modules are enriched with genes which are involved with extracellular matrix structural components (bisque4) and DNA replication and repair and the cell cycle (turquoise). These analyses further support our original observation that indicate unique microglia transcriptional responses to different species of Aβ peptides.

Interestingly, sub-populations of microglia previously identified in sc-, sn-RNA-seq or spatial transcriptomic studies did not fall within any single module (Figure 4B). For example, plaque-induced genes (PIGs) which are found in microglia surrounding Aβ plaques (Chen *et al*., 2020) fall across multiple modules and those modules do not fit any pattern of being correlated or not with any treatment paradigm including being both negatively and positively correlated with various treatments. This pattern also holds for genes found within the neurodegenerative disease-associated phagocytic microglia cells (DAMs) (Keren-Shaul *et al*., 2017) as well as microglia associated with a neurodegenerative phenotype (MGnD) (Krasemann *et al*., 2017). As noted in these prior studies, the microglia sub-populations share a number of genes in common.

To examine the strength of gene-gene connections with these networks, we chose representative modules that were positively correlated in only one treatment type and examined the networks across all treatments (Figure 5). We plotted edge weights to represent gene-gene connection strengths in an ordered heatmap to visualize the overall network strength more easily between the various treatment paradigms. Given that the salmon module has the strongest correlation value with the 1-hr fAβ treatment, we used this as a representative network for acute, 1-hr fAβ treatment. We find the genes within the salmon module have a stronger overall connection as compared with long-term, 12-hr fAβ or oAβ treatments (Figure 5A). Similar enhancements in gene-gene network strength were seen for the blue module which is positively correlated with long-term, 12-hr fAβ treatment (Figure 5B). Though this module does not have the strongest correlation value of the five modules highly correlated with the 12- hr fAβ treatments, we chose to examine this module as its member genes are enriched in interferon and immune signaling pathways. Finally, the turquoise module, which has the highest correlation value of the five modules highly correlated with oAβ treatment, was chosen as the representative module for positive correlation with long-term, 12-hr oAβ treatment (Figure 5C). This module also shows a striking increase in network connection strength as compared with the other two conditions. Genes within this module show enrichment in cell cycle, DNA replication and repair pathways. The top hub genes for these three modules are listed in Table 2.

**Figure 5:**
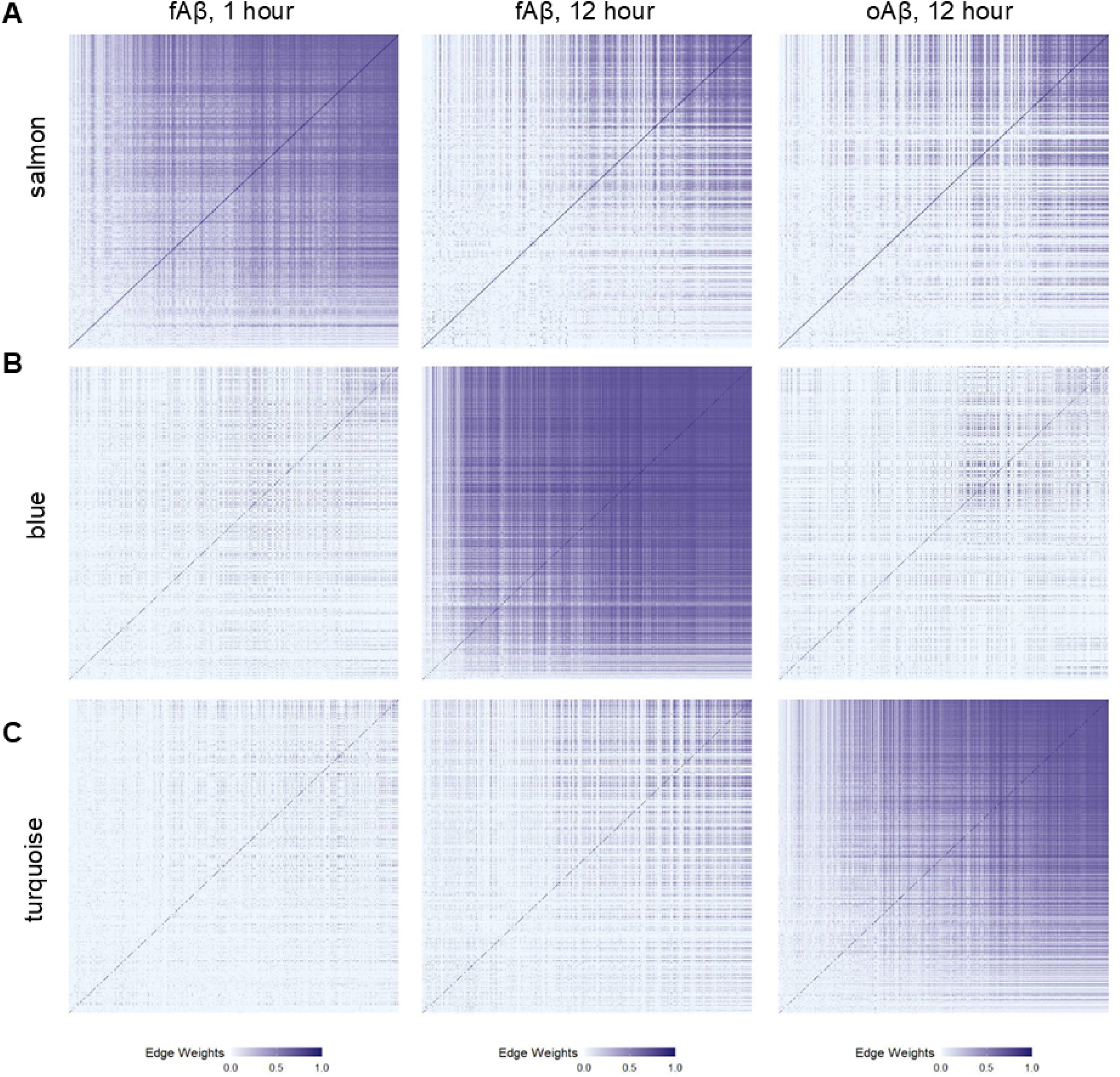
Gene networks are strongest in the modules that are positively correlated with Aβ treatments. Gene networks are shown as heatmaps of the edge weight. A greater edge weight (darker blue shades) indicates a strong gene-gene connection. The order of genes within each heatmap is preserved for the comparisons across Aβ treatment types. A) The gene network for the salmon module, a representative module positively correlated with acute, 1-hr fAβ treatment, is strongest than in 12-hr fAβ or 12-hr oAβ treatments. B) The gene network for the blue module, a representative module positively correlated with long-term, 12hr fAβ treatment is stronger than in 1-hr fAβ or 12-hr oAβ conditions. C) The gene network for the turquoise modules, which represent a module positively correlated with long-term, 12- hr oAβ treatment is strongest in oAβ treatments.

**Table 2:** WGCNA module statistics. Top 10 hub genes within the modules depicted in Figure 5 following WGCNA. Results are sorted by Gene Significance to each treatment type. Module statistics including gene significance value (to treatment), p-values corresponding to the gene significance (GS p-value), module membership value (of gene to module), module membership p-value (MM p-value) and gene connectivity within the module (kWithin) are shown.

| Hub Gene | Gene Significance (GS) | GS p-value | Module Membership (MM) | MM p-value | kWithin |
| --- | --- | --- | --- | --- | --- |
| **salmon** | to fAβ42, 1-hr | | | | |
| **Gabbr2** | 0.9984 | 8.34E-14 | 0.9854 | 5.17E-09 | 167.213 |
| **Vegfa** | 0.9980 | 2.47E-13 | 0.9927 | 1.58E-10 | 166.033 |
| **Cyth1** | 0.9977 | 4.60E-13 | 0.9953 | 1.79E-11 | 164.974 |
| **Rab7b** | 0.9976 | 5.71E-13 | 0.9903 | 6.51E-10 | 159.731 |
| **Tnfrsf21** | 0.9971 | 1.72E-12 | 0.9929 | 1.42E-10 | 164.391 |
| **Gpcpd1** | 0.9965 | 3.83E-12 | 0.9920 | 2.47E-10 | 157.545 |
| **Usp2** | 0.9962 | 6.38E-12 | 0.9498 | 2.32E-06 | 155.563 |
| **Folr2** | 0.9961 | 6.94E-12 | 0.9800 | 2.46E-08 | 167.461 |
| **Picalm** | 0.9961 | 7.20E-12 | 0.9726 | 1.16E-07 | 159.591 |
| **Plek** | 0.9959 | 8.80E-12 | 0.9896 | 9.34E-10 | 159.217 |
| **blue** | to fAβ42, 12-hr | | | | |
| **Tmem176a** | 0.9997 | 3.13E-17 | 0.9513 | 1.98E-06 | 468.954 |
| **Acp2** | 0.9996 | 4.65E-17 | 0.9321 | 1.02E-05 | 462.765 |
| **Gpr18** | 0.9996 | 5.40E-17 | 0.9432 | 4.24E-06 | 450.890 |
| **Fnbp1l** | 0.9995 | 1.77E-16 | 0.9736 | 9.61E-08 | 452.921 |
| **Tmem176b** | 0.9995 | 2.50E-16 | 0.9316 | 1.05E-05 | 479.564 |
| **Adgre1** | 0.9994 | 5.13E-16 | 0.9676 | 2.65E-07 | 485.170 |
| **Slc11a2** | 0.9994 | 5.51E-16 | 0.9240 | 1.76E-05 | 487.036 |
| **Cep85** | 0.9991 | 5.73E-15 | 0.9440 | 3.94E-06 | 463.008 |
| **Cd82** | 0.9989 | 1.12E-14 | 0.9493 | 2.43E-06 | 462.487 |
| **Nr1h3** | 0.9989 | 1.34E-14 | 0.9719 | 1.33E-07 | 466.064 |
| **turquoise** | to oAβ42, 12-hr | | | | |
| **Asf1b** | 0.9998 | 6.09E-19 | 0.991 | 3.71E-10 | 400.289 |
| **Alox5** | 0.9998 | 2.49E-18 | 0.933 | 9.27E-06 | 402.563 |
| **S100a4** | 0.9998 | 3.79E-18 | 0.973 | 1.18E-07 | 400.107 |
| **Klf2** | 0.9996 | 4.53E-17 | 0.952 | 1.89E-06 | 402.789 |
| **Hal** | 0.9996 | 7.56E-17 | 0.868 | 2.49E-04 | 401.621 |
| **Top2a** | 0.9996 | 7.82E-17 | 0.996 | 6.24E-12 | 401.975 |
| **Cks1b** | 0.9996 | 8.13E-17 | 0.950 | 2.16E-06 | 403.304 |
| **Pygl** | 0.9991 | 3.81E-15 | 0.961 | 6.31E-07 | 403.646 |
| **E2f1** | 0.9991 | 5.78E-15 | 0.965 | 3.66E-07 | 402.021 |
| **Mcm7** | 0.9989 | 1.35E-14 | 0.958 | 9.81E-07 | 400.930 |

### Transcriptional changes in primary microglia do not mimic those seen in the transgenic CRND8 mouse brain

To understand how well *ex vivo* changes in primary microglia cultures recapitulate *in vivo* processes, we examined transcriptional changes in the brains of transgenic amyloid mouse model CRND8 at 3, 6, 12 and 20 months of age by bulk RNA-seq. Using the same cut-off values to identify differentially expressed genes as above, we find that at 3 months of age there are few transcriptional changes between the transgenic CRND8 and their non-transgenic littermate controls (11 upregulated, 4 downregulated, Figure 6A, Supplemental Data 12). By 6 months of age, the number of transcriptional changes increases to 187 upregulated and 105 downregulated genes (Figure 6B). At 12 months of age, the number of differentially expressed genes is higher than at previous timepoints and is dominated by changes in upregulated genes (493 upregulated genes) over those that are downregulated (103 downregulated genes, Figure 6C, Supplemental Data 14). At 20 months, more genes continue to be upregulated (746 genes) than downregulated (115 genes, Figure 6D, Supplemental Data 15). Trends for GO term enrichment in downregulated genes was not evident until 12 months with enriched terms including a variety of receptor binding activities (i.e., *glucocorticoid receptor binding, steroid hormone receptor* activity) and involvement of core promoter activity (*core promoter sequence-specific DNA binding*; Figure 6E). Upregulated genes are enriched primarily with immune responses (*immunoglobulin receptor activity, IgG binding*) that are consistent as the mice age (Figure 6F, Supplemental Data 13).

**Figure 6:**
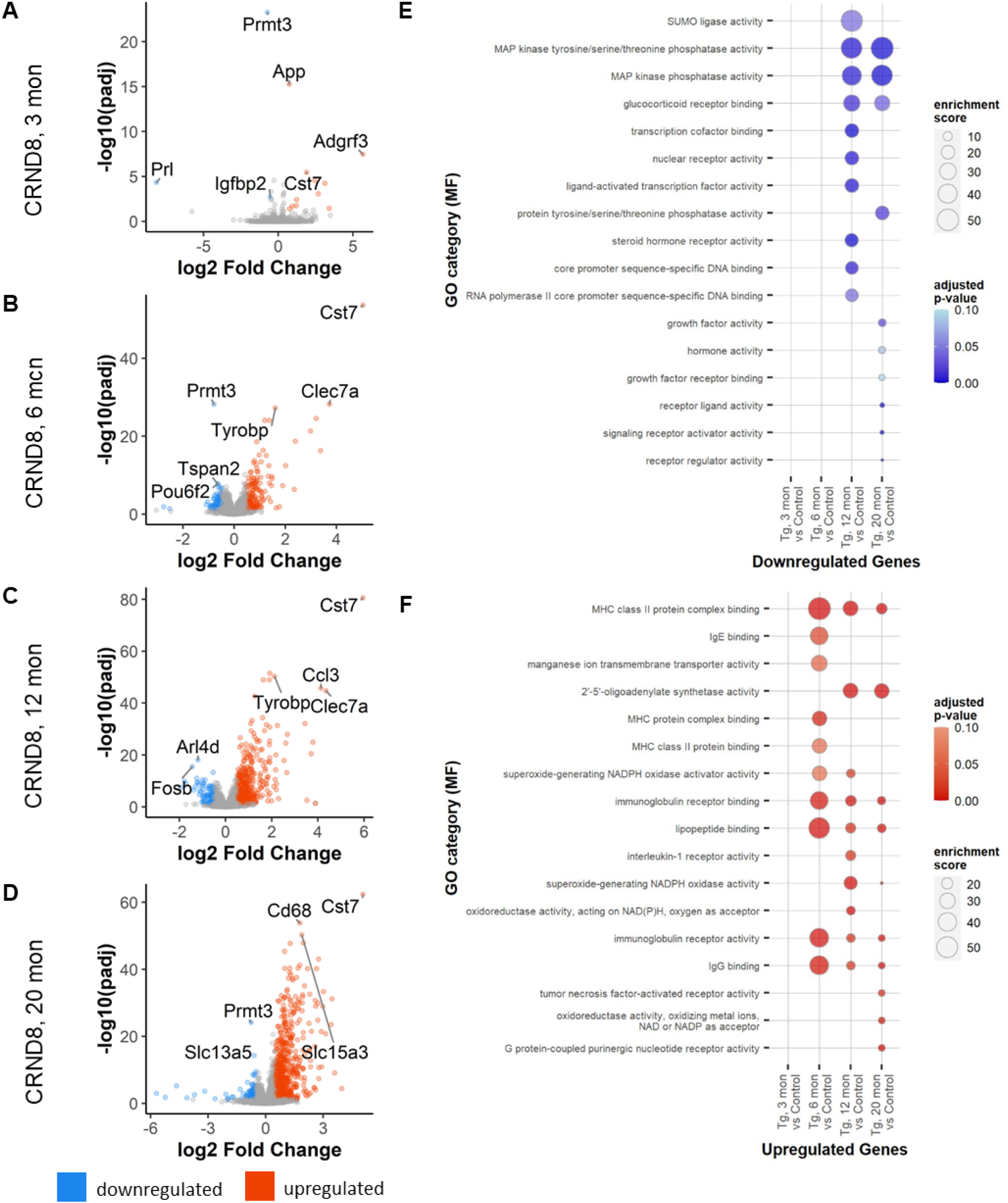
Differential gene expression in transgenic CRND8 mice. A) Total changes in down- (blue) and up- (red) regulated genes in transgnic CRND8 mouse brains versus non-transgenic controls at 3 months. B) Total changes in down- and upregulated genes in transgenic CRND8 mouse brains versus non-transgenic controls at 6 months. C) Total changes in down- and upregulated genes in transgenic CRND8 mouse brains versus non-transgenic controls at 12 months. D) Total changes in down- and upregulated genes in transgenic CRND8 mouse brains versus non-transgenic controls at 20 months. E) Bubble plots of GO category enrichment results for downregulated genes. F) Bubble plots of GO category enrichment results for upregulated genes. Plots for GO category over-enrichment analysis show the top 10 hits for each comparison by enrichment score following a filter step by a p-value adjusted for multiple comparisons of ≤ 0.1 and keeping GO categories with greater than 5 genes within the category.

Not surprisingly, direct comparisons of the microglial-specific genes in Aβ-treated primary microglia with transgenic CRND8 mice are poorly correlated (Figure 7). Correlation values are low between either differentially expressed microglial genes in transgenic CRND8 mice at 20 months versus Aβ-treated primary microglia for any treatment paradigm, oAβ 12-hr (Figure 7A), fAβ 12-hr (Supplemental Figure 3A) or fAβ 1-hr (Supplemental Figure 3B). In the transgenic CRND8, these genes are nearly universally upregulated, but are both up- and down-regulated in the primary microglia. We examined representative genes that are highly differentially expressed in Aβ-treated microglia (Figure 7D) which reveal little (Vim) to no (Sod2, Sgk1) corresponding changes in the transgenic CRND8 mice over time—indeed some changes were opposite of those observed in CRND8 (Fcgr2b). Conversely, examining a selection of genes that are consistently and significantly changed in transgenic CRND8 mice over time (Figure 7E; Cst7, Irf8 and Plek) exposes variable responses in microglia after the application of either fAβ or oAβ peptide. Additionally, a panel of Alzheimer’s-disease relevant genes, which are consistently upregulated in the transgenic CRND8 mouse brain over time, also reveals variable (Abi3 vs Plcg2)—and sometimes unexpected (Trem2)—responses to Aβ peptides in microglia (Figure 7F). This pattern is also seen in a selection of cytokines (Figure 7G; Ccl3, Ccl4 and Tnf) and cytokine receptor (Figure 7H; Ccr5, Csf3r and Tnfrsf1a) genes.

**Figure 7:**
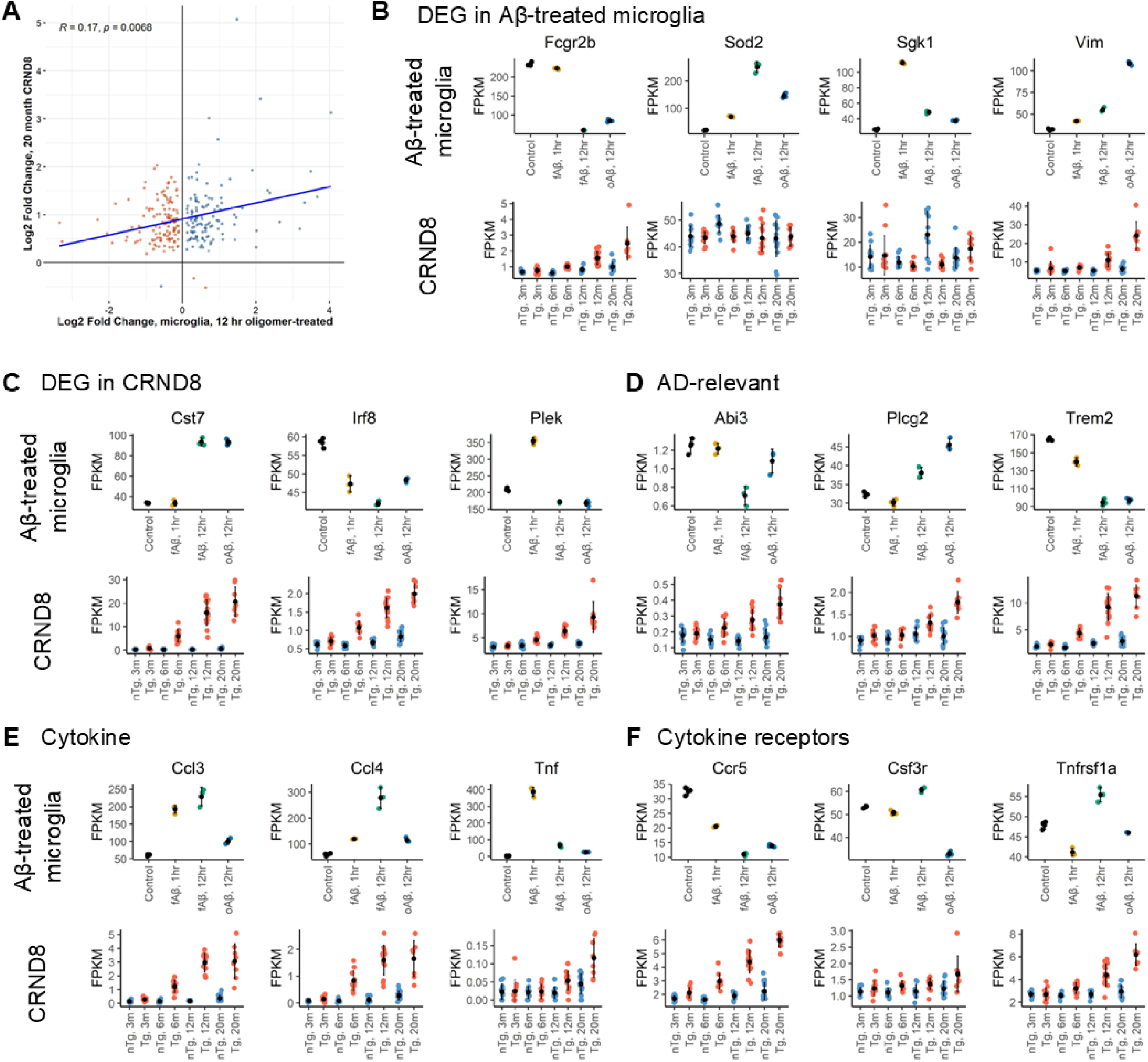
Microglia transcriptional responses at the individual gene level are not reflective of changes seen in the CRND8 model. A) Comparisons of log_2_ fold change values for microglial genes (Zhang *et al*., 2014) in transgenic CRND8 versus oAβ-treatment in primary microglia show little correlation. Geometric means of FPKM data of representative genes differentially expressed in Ab-treated primary microglia is shown for Aβ-treated microglia (top row) and CRND8 mouse brains (bottom row) (B). Similar plots are shown for representative differentially expressed genes identified in CRND8 mice (C), AD-relevant genes (D), representative cytokine genes (E) and representative cytokine receptor genes (F).

We then examined the transcriptional profile of microglial cell subsets that have been identified in past sc-, sn-RNA-seq or spatial transcriptomic studies of microglia (Figure 8). As evidenced by the increase in the transgenic CRND8 brains, the expression of these genes within these subpopulations is increased in AD. A general transcriptomic signature of microglial-enriched genes (Zhang *et al*, 2014) is increased following all Aβ treatments in microglia primary cultures—a signal that mimics increases seen in transgenic CRND8 mice over time (Figure 8A). A homeostatic microglia (H2M) signature (Sala Frigerio *et al*, 2019) increases over time in transgenic CRND8 mice, but this increase is seen only in microglial cultures treated for 12-hr fAβ (Figure 8B). We examined the transcriptional signature associated with cycling and proliferative microglia (Sala Frigerio *et al*., 2019) (Figure 8C). There is a large increase in the CPM signature in oAβ treated microglia, but no difference is seen in the transgenic CRND8—which stands as a contrast to the general trend in the other microglial subtypes. This likely reflects that this population represents a very small percentage of microglial cells within the brain (Sala Frigerio *et al*., 2019) and its signature is lost within the larger milieu of other cell types within the brain. We also examine interferon-responsive microglia (IRMs; Figure 8D) (Sala Frigerio *et al*., 2019). This transcriptional signature increased over time in transgenic CRND8 but a large change is seen only in response to long-term fAβ treatment. Interestingly, an increased transcriptional response in the disease associated microglia profile (DAMs, found in the microglia surrounding Aβ plaques (Keren-Shaul *et al*., 2017)) is seen in response to fAβ, but not oAβ treatment, while a steady increase is seen in the transgenic CRND8 (Figure 8E). Intriguingly, transcriptional responses linked to both activated response microglia (ARMs, (Sala Frigerio *et al*., 2019) which are responsive to Aβ deposition) as well are plaque-induced genes (PIGs; (Chen *et al*., 2020) are decreased or unchanged in all treatment paradigms in primary microglia while these genes steadily increase over time in transgenic CRND8. Genes linked to the microglial neurodegenerative phenotype (MGnD) (Krasemann *et al*., 2017) appear as a likely reliable indicator of transcriptional changes for all Aβ treatment paradigms as well as in transgenic CRND8 brains.

**Figure 8:**
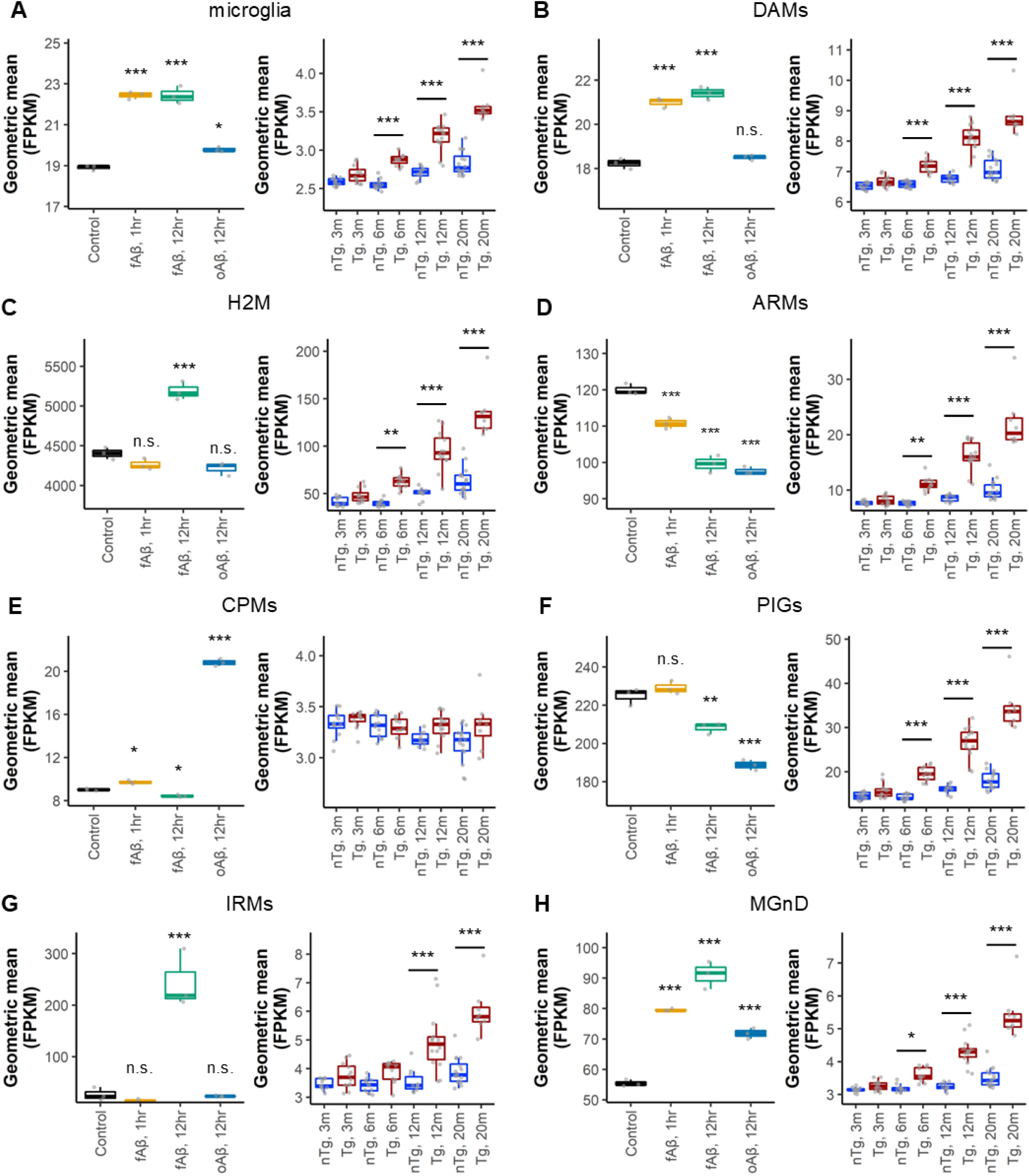
Microglia sub-type transcriptional signatures in primary microglia do not reflect changes seen in the CRND8 model. Gene signatures for microglia genes and sub-populations of microglia are shown for primary microglia cultures (left) and CRND8 mouse brains (right). A) Microglia expression signature identified in Zhang *et al*., 2014. B) Activated microglia expression signature in Aβ-treated microglia. B) Disease-associated microglia (DAM) gene expression signature identified in Keren Shaul *et al.,* 2017. C) Homeostatic (H2M) microglial gene expression signature as in Sala Frigerio *et al*., 2016. D) Activated response microglia (ARM) gene expression signature as in Sala Frigerio *et al*., 2016. E) Cycling and proliferating microglia (CPM) gene expression signature as in Sala Frigerio *et al*., 2016. F) Plaque-induced microglia (PIG) gene expression signature as in Chen *et al*., 2020. G) Interferon-responsive microglia (IRM) gene expression signature as in Sala Frigerio *et al*., 2016. H) Neurodegenerative microglia phenotype (MGnD) gene expression signature as in Karesmann, *et al*., 2017. *, p-adj < 0.05; **, p-adj < 0.01; ***, p-adj < 0.001.

## Discussion

Acute exposure of cultured primary microglia to oAβ or fAβ elicits a robust and rapid transcriptional response. Both forms of Aβ induce significant increases and decreases in RNA levels for hundreds of genes. Nevertheless, transcriptomic responses to oAβ and fAβ at 12-hr are distinguishable. Of note, the finding that oAβ increases RNAs associated primarily with cell cycle whereas fAβ increases RNAs associated primarily with phagocytic processes is intriguing.

As there are numerous validated and candidate Aβ receptors expressed on microglia (Jarosz-Griffiths *et al*, 2016), such studies indicate that acute exposure to Aβ aggregates induces robust cellular events that can be assessed at the systems level using transcriptomic approaches. Based on the studies of fAβ there is a clear temporality to the response with varying clusters of genes changing in both similar and different directions at the various time points. These data are reminiscent of studies examining acute effects of LPS on primary microglia, though given numerous experimental differences with historical data sets a much more systematic, side by side, comparison would be needed to evaluate the overall similarity in response to classic proinflammatory mediators such as LPS and Aβ.

As we and others have used primary microglia to study uptake and clearance of Aβ and Aβ aggregates, a primary objective of this study was to determine if the response to Aβ in such acute studies is indicative of system levels changes in mouse models of Aβ deposition, where Aβ accumulates over time. In this case, we have compared the transcriptomic changes in CRND8 transgenic model (compared to non-transgenic controls) with our acute transcriptomic signatures of the primary microglia exposed to Aβ. These data reveal that acute transcriptional responses of primary microglia to Aβ poorly reflect the *in vivo* responses of genes to chronic progressive Aβ accumulation. Like a number of other recent studies, these data suggest that though primary microglial studies may have utility in some settings, extrapolating results from these studies to the *in vivo* setting is problematic.

Many laboratories in the field, including our own, have focused on responses of microglia to classic cytokines including but not limited to TNFα, IL1α, IL1β, IL10, IL6 and IFNγ (Chakrabarty *et al*, 2010; Chakrabarty *et al*, 2011; Chakrabarty *et al*, 2015; Colon-Perez *et al*, 2019; Webers *et al*, 2020). Though these cytokines show massive changes in transcript in primary culture, *in vivo* transcript levels in the brain are very low throughout the lifespan of the non-Tg and Tg mice. Though some cytokines show small increases over time in the presence of amyloid deposition, the magnitude of this increase is nowhere near the scale of increase observed in the primary culture. The massive increases in transcript levels observed in primary microglial cultures of many of these cytokines and other immune factors have likely contributed to the field’s focus on these as key mediators of the microglial responses to Aβ and other insults. However, data presented here, as well as other studies (Butovsky *et al*., 2014), highlight differences between the *ex vivo* and *in vivo* microglial responses and indicate that the focus on some of these cytokine and other immune factors may be misleading.

Notably, microglia—at least at the transcript level—express moderate to high levels of many classic cytokine receptors *in vivo*. Perhaps, the low level of ligand expression compared to relatively high levels of receptor would suggest that these receptors on microglia serve primarily to sense non-CNS changes in the cytokine levels following peripheral insults. In any case, these data along with numerous other studies demonstrating the heterogeneity of microglia *in vivo* (Butovsky *et al*., 2014; Chen *et al*., 2020; Friedman *et al*., 2018; Hammond *et al*., 2019; Keren-Shaul *et al*., 2017; Krasemann *et al*., 2017; Olah *et al*., 2020), highlight the notion that primary isolated microglia cells are poor proxies for *in vivo* responses. As study of microglial cells in the brain has many limitations, additional efforts to develop better *ex vivo* models of microglial responses would benefit the field. Although several reports of such efforts exist (Arber *et al*, 2017; Croft *et al*, 2019), further evaluation and “stress-testing” of these and other *ex vivo* methods will be needed before they are likely to be widely adopted.

Previous studies have focused on the functional consequences of treating various primary CNS cells with oAβ or fAβ. oAβ species have been conceptualized by some in the field as the proximal neurotoxin in AD (Cline *et al*, 2018; Haass & Selkoe, 2007; Li & Selkoe, 2020; Wang *et al*., 2016), as they disrupt synaptic transmission in neurons at very low, picomolar concentrations (Rammes *et al*, 2011; Waters, 2010). However, the evidence that oAβ species are overtly toxic with respect to inducing neuronal death is lacking; further there is debate as to whether appreciable concentrations of intrinsically soluble oligomers exists in the AD brain or mouse models of amyloid deposition (Jan *et al*, 2011; Jan *et al*, 2008; Tseng *et al*, 1999; van Helmond *et al*, 2010). In contrast, at least in primary neuronal cultures higher concentrations of various aggregates have been linked to induction of neuronal death via apoptotic mechanisms (Deshpande *et al*, 2006). Both direct toxicity of the aggregates, or aggregate growth and indirect toxicity via activation of glial cells that results in neurotoxicity have been invoked as mechanisms underlying Aβ induced neuronal death (Kayed & Lasagna-Reeves, 2013). Clearly, the massive alterations in microglial cells observed here in response to synthetic Aβ aggregates reinforces the potential for neurotoxicity in mixed primary cultures. However, we would note that both oAβ and fAβ induce massive changes in the transcriptome of microglia, and certainly lend little credence to claims by some in the field that fAβ is inert. Indeed, our results suggest that microglial transcriptional responses to fAβ more closely mimic *in vivo* responses to amyloid accumulation as evidenced by the behavior of the “blue” module genes in our study which positively correlated with fAβ treatment are paralleled in the transgenic CRND8 brain and the microglial sub-type analysis.

As suggested above, the concept that microglial cells might make exquisitely sensitive biosensors that can be used to distinguish between various aggregate forms is intriguing. Microglial do appear at the transcript level to respond in partially overlapping, but distinct ways to oAβ or fAβ. Much more extensive studies will be needed to follow up on this intriguing observation. However, from a teleological point of view this concept makes quite a bit of sense. Microglial cells with a plethora of damage associated and pathogen associated receptors are designed to respond rapidly to potentially harmful proteins and other stimuli (Deshpande *et al*., 2006). One would predict that overlapping but distinct binding interactions could result in partially overlapping but distinct responses that might essentially provide a type of integration of signals to distinguish various aggregates.

As the main goal of these studies was to assess the system level responses of microglial cells in culture to Aβ aggregates and compare that to a longitudinal transcriptomic study in APP mice, there are a number of limitations that are worth noting. First, both dose response and more extended time courses were not conducted. Second, we did not include monomeric Aβ42, as it would likely aggregate at these concentrations during incubation; nor did we include a short-term oAβ42 timepoint. Third, we did not extensively purify oligomeric assemblies to a more defined species. Finally, we have not pursued studies to determine whether fAβ and oAβ induce different functional states in the cultured microglia cells. It is almost certain such studies would yield interesting data, but it is unlikely that it would alter the relevance of the work with respect to disease implications in AD.

A recent elegant study exploring *in vivo* microglial responses to LPS using translational profiling approaches to assess both ribosome-associated transcripts and proteins, showed major discrepancies between the proinflammatory transcriptomic signature and a more immune modulatory and homeostatic protein signature (Boutej *et al*, 2017). Given the massive upregulation of proinflammatory transcripts in cultured microglia exposed to Aβ and the large number of upregulated microglial transcripts in APP mouse models and human AD, it will be important to integrate proteomic and transcriptomic studies of microglia in the future. Indeed, at least in the Boutej study, the biologic inferences derived from evaluating the proteome or transcriptome are disparate and only when the two are compared directly does the concept of widespread translation repression emerge. Additional studies also show that even the process of rapid isolation of microglial cells from the brain changes their transcriptome (He *et al*, 2018; Lin *et al*, 2017; Tham *et al*, 2003). Thus, even though single cell transcriptomic and proteomic studies of isolated microglia cells potentially provide new insights into their roles in health and disease, additional validation using *in situ* methodologies is needed to confirm that changes observed reflect changes *in situ* and are not induced during the isolation.

The number of studies focusing on microglia cells and their impact on AD and other neurodegenerative disorders is rapidly expanding. This study and many others highlight that traditional methods to study them, such as in primary cultures, are highly artificial and may lead to inappropriate conclusions. Current efforts to develop strategies to harness microglial function in a therapeutically beneficial fashion, must by necessity study the effect of that therapy *in vivo*. However, given the large number of immune factors that are emerging as modulators of neurodegenerative pathologies, and the limitations of only studying these cells *in vivo*, additional efforts to validate *ex vivo* systems that better approximate microglial functions *in vivo* ware warranted.

## Figure Legends

**Supplemental Figure 1: Comparison of gene expression changes in Aβ42-treated microglia.**

Comparisons of significantly changed genes between each Ab treatment paradigm are plotted by log_2_ fold change. Genes with congruent changes (blue) are in the upper, right (commonly upregulated) and lower, left (commonly downregulated) quadrants. Disparate changes in gene expression (orange) are seen in the upper, left quadrant (upregulated in the first but downregulated in the second condition) and in the lower, right quadrant (downregulated in the first but upregulated in the second condition). A Spearman’s correlation analysis was performed, and results are indicated by the blue line with R and p-values as indicated. A) oAβ-treated microglia at 12-hr (versus control) compared against changes seen in fAβ-treated microglia at 12-hr (versus control). B) Bubble plot of GO category over-enrichment analysis of genes in each plot quadrant in A. C) oAβ-treated microglia at 12-hr (versus control) compared with changes seen in fAβ-treated microglia at 1-hr (versus control). D) Bubble plot of GO category over-enrichment analysis of genes in each plot quadrant in C. E) fAβ-treated microglia at 12-hr (versus control) compared with changes seen in fAβ-treated microglia at 1-hr (versus control). G) Bubble plot of GO category over-enrichment analysis results for genes in all four plot quadrants in E. Plots of GO category over-enrichment analysis show the top 10 categories by enrichment score following a filtering step by a p-value adjusted for multiple comparisons testing of ≤ 0.1 and removing GO categories with less than 5 genes within the category.

## Methods

### Animal Research

All animal research was performed under protocols approved by the Institute for Animal Care and Use Committee (IACUC) at the University of Florida.

### Microglial primary cultures and Aβ42 treatment

Mouse pups for primary microglial cultures are obtained from matings of B6/C3HF1 mice (Envigo). Mice are given ad libitum access to food and water and are maintained on a 12-hr light/12-hr dark cycle. Primary microglia cultures were isolated following described protocols (Rosario *et al*, 2016). Briefly, cortices were isolated at post-natal day P2-P3. The mixed microglial/astrocyte cultures were maintained in 75cm^2^ flasks with 20 ml of DMEM containing 10% fetal bovine serum. After 10 days, the flasks were shaken for 30 minutes at 37°C at 150 rpm to dislodge the microglia from the adherent astrocyte layer. The microglia were plated into 6-well plates and maintained at 37°C. One day after plating, microglia were treated with 5 μM Aβ_42_ fibrils or oligomers for 1- or 12-hr as noted. Cell were washed with PBS prior to harvest. Three replicates for each condition were done.

### Fibrillar and oligomeric Aβ preparation

Fibrillar and oligomeric forms of Aβ were prepared as previously described (Chakrabarty *et al*, 2018; Stine *et al*, 2003). Aliquots (10, 100 and 1000 ng) were separated on SDS-PAGE page run using Biorad Criterion 10% bis-tris gel and XT running buffer/sample buffer for 60m at 180V (constant). Gel transferred onto 0.2micron PVDF in Towbin transfer buffer for 45m at 150V (constant). 6E10 (Biolegend, San Diego, CA) primary antibody diluted at 1:1000 and applied for 1.5h at 37°. Primary antibody was detected with goat anti-mouse IR700 and scanned on LiCor Odyssey 700mm channel.

### RNA extraction and sequencing

Microglial RNA was extracted using the RNeasy mini extraction kit with on-column DNase treatment (Qiagen). RNA quality was determined with the Qubit RNA HS assay. RNA quality was checked via RIN on an Agilent Bioanalyzer 2100 with the Eukaryote Total RNA Nano chip. Libraries were generated with the Illumina RNA-seq library prep for low input RNA. Libraries were sequenced on paired-end, 75 bp runs on the Nextseq 500 (Illumina). RNA QC, library preparation and sequencing were performed at the University of Florida’s Interdisciplinary Center for Biotechnology Research (ICBR) sequencing core.

### Transgenic CRND8 RNA-sequencing data

Data for the transgenic CRND8 mice was obtained from the AMP-AD Knowledge Portal (doi: <u>10.7303/syn3157182</u>). Experimental details are located within the data portal’s website. BAM files were downloaded from the AD Knowledge portal and used with the analysis method described below. Animal numbers are as follows: 3-month, nTg-F: 6; 3-month, nTg-M: 6; 3-month, Tg-F: 6; 3- month, Tg-M: 6; 6-month, nTg-F: 5; 6-month, nTg-M: 7; 6-month, Tg-F: 5; 6-month, Tg-M: 6; 12-month, nTg-F: 5; 12-month, nTg-M: 5; 12-month, Tg-F: 7; 12-month, Tg-M: 7; 20-month, nTg-F:11; 20-month, nTg-M: 5; 20-month, Tg-F: 5; 20-month, Tg-M: 3. Male and female mice of the same age and genotype were grouped together for this analysis.

### RNA-seq analysis

#### FASTQ alignment, gene counts and differential expression analysis

Resulting FASTQ files were aligned against the mouse genome (GRCm38) and GRCm38.94 annotation using STAR v2.6.1a (Dobin *et al*, 2013) to generate BAM files. BAM files were used to generate gene counts were generated using Rsamtools (Morgan *et al*, 2018) and the summarizeOverlaps function with the GenomicAlignments package (Lawrence *et al*, 2013). Differential gene expression analysis was performed with DESeq2 package using the “DESeq” function with default settings (Love *et al*, 2014) which fits a generalized linear model for each gene. Subsequent Wald test p-values are adjusted for multiple comparisons using the Benjamini-Hochberg method (adjusted p-value). Pair-wise changes in gene expression levels were examined between groups to identify differentially expressed genes (DEGs). DEGs were defined as an absolute log2Fold Change ≥ 0.5 and an adjusted p-value ≤0.05.

#### WGCNA

The WGCNA package in R (Langfelder & Horvath, 2008, 2012) was used to construct gene correlation networks from the expression data after filtering and removing genes with zero variance. For the microglia dataset, a soft power setting of 9 was chosen using the “pickSoftThreshold” function within the WGCNA package. The network was constructed using all microglial samples. Adjacency matrices were constructed using expression data and these power setting with the “adjacency” function and a signed hybrid network. Module identification was performed using the “cutreeDynamic” function and a deepSplit setting of 2 with a minimum module size of 30 for all analyses.

#### Functional annotation of DEGs, heatmap clusters and WGCNA modules

Gene ontology enrichment analysis was performed with goseq v1.42.0 (Young *et al*, 2010) to identify gene ontology categories—focusing on the molecular function (MF) category—and KEGG pathways that are affected between the various conditions. For DEGs, up- and down-regulated gene lists were analyzed separately. For WGCNA, gene lists from each module were used as input. Over-represented p-values were adjusted for multiple comparisons using the Benjamini-Hochberg adjustments for controlling false-discovery rates. An enrichment score was calculated by an observed-over-expected ratio of

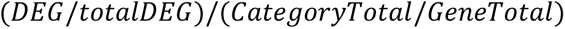

Where *DEG* represents the total number of DEGs or module genes within the GO or KEGG category, *totalDEG* represents the total number of DEGs or module genes; *CategoryTotal* represents the total number of genes within the GO or KEGG category and *GeneTotal* represents the total number of genes examined. GO terms and KEGG pathways are filtered for p-values adjusted for multiple comparisons (BHadjust) < 0.05 (Aβ-treated microglia) or 0.1 (CRND8 mice), enrichment scores > 1 and total number of genes within the category > 5.

Z-scores for genes identified as a DEG for any Aβ-treatment comparison versus control were plotted in a heatmap using pheatmap v1.0.12. Clusters were identified using the cutree function with h = 5.75. goseq was used for GO and KEGG pathway analysis on genes within each cluster. GO terms and KEGG pathways are filtered for p-values < 0.05, enrichment scores > 1 and total number of genes within the category > 5.

Gene lists to annotate WGCNA modules and identify microglia subtype signatures were identified from previously published studies (Chen *et al*., 2020; Friedman *et al*., 2018; Hammond *et al*., 2019; Keren-Shaul *et al*., 2017; Krasemann *et al*., 2017; Sala Frigerio *et al*., 2019; Zhang *et al*., 2014) (see also Supplemental Data 10). Gene overlap analysis was conducted with the GeneOverlap package in R (Shen, 2020). GeneOverlap uses Fisher’s exact test to calculate the p-value for significance testing as well as calculating the odds ratio. goseq was used for GO and KEGG pathway analysis of genes within each module filtering for those terms with p-values < 0.05, enrichment scores > 1 and total number of genes within the category > 5.

#### Direct comparisons of DEGs between treatment types

DEG datasets for each treatment paradigm against control were filtered for significant gene changes using criteria described above. The distribution of resulting log2FoldChange was tested for a normal distribution using the Shapiro-Wilk Normality Test. The correlation value for the log2FoldChange value in each pair-wise comparison was calculated using Spearman’s rank-order correlation test at a confidence level set to 0.95 in R and graphs were drawn using the ggpubr package in R (Kassambara, 2020).

#### Statistical analysis and data visualizations

ANOVA with Tukey’s post-hoc multiple comparisons test was performed in R. Data visualizations were generated in R using the ggplot2 package (Wickham, 2016) unless otherwise noted. Bar plots show mean ± SD. For boxplots, upper, middle and lower hinges correspond to first quartile, median and third quartiles, respectively. Upper (or lower) whiskers correspond to the largest (or smallest) observation beyond the upper hinge up to 1.5 times the inter-quartile range. Outliers beyond the upper and lower whiskers are plotted independently.

### Data Availability

FASTQ files for the Aβ-treated primary microglia samples are available via the AD Knowledge Portal (https://adknowledgeportal.org). The AD Knowledge Portal is a platform for accessing data, analyses, and tools generated by the Accelerating Medicines Partnership (AMP-AD) Target Discovery Program and other National Institute on Aging (NIA)-supported programs to enable open-science practices and accelerate translational learning. The data, analyses and tools are shared early in the research cycle without a publication embargo on secondary use. Data is available for general research use according to the following requirements for data access and data attribution (https://adknowledgeportal.org/DataAccess/Instructions).

For access to content described in this manuscript see: http://doi.org/10.7303/syn25006578

Interactive data portals are available for viewing at the following: Aβ-treated microglial DEG data: https://tinyurl.com/y3q3kaoe

CRND8 DEG data: https://tinyurl.com/y5evwkuw

cross-treatment comparisons of DEG data: https://tinyurl.com/yyph68vc

## Acknowledgements

Support for these studies was provided by the NIH/NIA U01 AG046139, P30 AG066506, P50 AG047266. NET is also funded in part by NIH/NIA RF1 AG051504, and R01 AG061796.

## Supplemental Data

### Supplemental Data 1

DEG results from primary microglia treated with fAβ, 1-hr vs Control.

Tab 1: Results from DESeq2 from the comparison of fAβ, 1-hr vs Control.

Tab 2: GOseq results from GO analysis on upregulated genes in fAβ, 1-hr vs Control. Categories are

Tab 3: GOseq results from GO analysis on downregulated genes in fAβ, 1-hr vs Control.

Tab 4: GOseq results from KEGG pathway analysis on upregulated genes in fAβ, 1-hr vs Control.

Tab 5: GOseq results from KEGG pathway analysis on downregulated genes in fAβ, 1-hr vs Control.

### Supplemental Data 2

DEG results from primary microglia treated with fAβ, 12-hr vs Control.

Tab 1: Results from DESeq2 from the comparison of fAβ, 12-hr vs Control.

Tab 2: GOseq results from GO analysis on upregulated genes in fAβ, 12-hr vs Control.

Tab 3: GOseq results from GO analysis on downregulated genes in fAβ, 12-hr vs Control.

Tab 4: GOseq results from KEGG pathway analysis on upregulated genes in fAβ, 12-hr vs Control.

Tab 5: GOseq results from KEGG pathway analysis on downregulated genes in fAβ, 12-hr vs Control.

### Supplemental Data 3

DEG results from primary microglia treated with oAβ, 12-hr vs Control.

Tab 1: Results from DESeq2 from the comparison of oAβ, 12-hr vs Control.

Tab 2: GOseq results from GO analysis on upregulated genes in oAβ, 12-hr vs Control.

Tab 3: GOseq results from GO analysis on downregulated genes in oAβ, 12-hr vs Control.

Tab 4: GOseq results from KEGG pathway analysis on upregulated genes in oAβ, 12-hr vs Control.

Tab 5: GOseq results from KEGG pathway analysis on downregulated genes in oAβ, 12-hr vs Control.

### Supplemental Data 4

DEG results from primary microglia treated with fAβ, 12-hr vs oAβ, 12-hr.

Tab 1: Results from DESeq2 from the comparison of fAβ, 12-hr vs oAβ, 12-hr.

Tab 2: GOseq results from GO analysis on upregulated genes in fAβ, 12-hr vs oAβ, 12-hr.

Tab 3: GOseq results from GO analysis on downregulated genes in fAβ, 12-hr vs oAβ, 12-hr.

Tab 4: GOseq results from KEGG pathway analysis on upregulated genes in fAβ, 12-hr vs oAβ, 12-hr.

Tab 5: GOseq results from KEGG pathway analysis on downregulated genes in fAβ, 12-hr vs oAβ, 12-hr.

### Supplemental Data 5

Cross-treatment comparison of DEGs in fAβ, 12h-hr (vs control) against oAβ, 12-hr (vs control).

Tab 1: GOseq results from GO analysis on genes found in each graph quadrant in Supplemental Figure 1A.

Tab 2: GOseq results from KEGG pathway on genes found in each graph quadrant in Supplemental Figure 1A.

### Supplemental Data 6

Cross-treatment comparison of DEGs in oAβ, 12h-hr (vs control) against fAβ, 1-hr (vs control).

Tab 1: GOseq results from GO analysis on genes found in each graph quadrant in Supplemental Figure 1B.

Tab 2: GOseq results from KEGG pathway on genes found in each graph quadrant in Supplemental Figure 1B.

### Supplemental Data 7

Cross-treatment comparison of DEGs in fAβ, 12h-hr (vs control) against fAβ, 1-hr (vs control).

Tab 1: GOseq results from GO analysis on genes found in each graph quadrant in Supplemental Figure 1C.

Tab 2: GOseq results from KEGG pathway on genes found in each graph quadrant in Supplemental Figure 1C.

### Supplemental Data 8

Analysis of gene clusters in Figure 3.

Tab 1: List of genes identified in each cluster. Genes that are significant in any treatment comparison versus control were plotted.

Tab 2: GOseq results from GO analysis on genes found within each cluster.

Tab 3: GOseq results from KEGG pathway analysis on genes found within teach cluster

### Supplemental Data 9

WGCNA results.

Tab 1: Gene-Module membership following WGCNA.

Tab 2: GOseq GO category analysis of genes with each module.

Tab 3: GOseq KEGG pathway analysis of genes within each module.

### Supplemental Data 10

Correlation of WGCNA modules with treatment paradigms

Correlation and associated p-values for relationships between WGCNA modules and Aβ treatments. Also included are the number of genes within each module and the top hub gene as identified by the “chooseTopHubInEachModule” function within the WGCNA package.

### Supplemental Data 11

Listing of published studies of single-cell, single-nuclear RNA-seq or spatial transcriptomic studies.

### Supplemental Data 12

DEG results from CRND8 mice, transgenic (Tg) versus non-transgenic (nTg), at 3 months.

Tab 1: DESeq2 results from transgenic (Tg) versus non-transgenic (nTg) mice at 3 months.

Tab 2: GOseq results from GO analysis on upregulated genes in Tg versus nTg, 3 months.

Tab 3: GOseq results from GO analysis on downregulated genes in Tg versus nTg, 3 months.

Tab 4: GOseq results from KEGG pathway analysis on upregulated genes in Tg versus nTg, 3 months.

Tab 5: GOseq results from KEGG pathway analysis on downregulated genes in Tg versus nTg, 3 months.

### Supplemental Data 13

DEG results from CRND8 mice, Tg versus nTg, at 6 months.

Tab 1: DESeq2 results from transgenic (Tg) versus non-transgenic (nTg) mice at 6 months.

Tab 2: GOseq results from GO analysis on upregulated genes in Tg versus nTg, 6 months.

Tab 3: GOseq results from GO analysis on downregulated genes in Tg versus nTg, 6 months.

Tab 4: GOseq results from KEGG pathway analysis on upregulated genes in Tg versus nTg, 6 months.

Tab 5: GOseq results from KEGG pathway analysis on downregulated genes in Tg versus nTg, 6 months.

### Supplemental Data 14

DEG results from CRND8 mice, Tg versus nTg, at 12 months.

Tab 1: DESeq2 results from transgenic (Tg) versus non-transgenic (nTg) mice at 12 months.

Tab 2: GOseq results from GO analysis on upregulated genes in Tg versus nTg, 12 months.

Tab 3: GOseq results from GO analysis on downregulated genes in Tg versus nTg, 12 months.

Tab 4: GOseq results from KEGG pathway analysis on upregulated genes in Tg versus nTg, 12 months.

Tab 5: GOseq results from KEGG pathway analysis on downregulated genes in Tg versus nTg, 12 months.

### Supplemental Data 15

DEG results from CRND8 mice, Tg versus nTg, at 20 months.

Tab 1: DESeq2 results from transgenic (Tg) versus non-transgenic (nTg) mice at 20 months.

Tab 2: GOseq results from GO analysis on upregulated genes in Tg versus nTg, 20 months.

Tab 3: GOseq results from GO analysis on downregulated genes in Tg versus nTg, 20 months.

Tab 4: GOseq results from KEGG pathway analysis on upregulated genes in Tg versus nTg, 20 months.

Tab 5: GOseq results from KEGG pathway analysis on downregulated genes in Tg versus nTg, 20 months.

## Notes

The authors declare they have no conflict of interest.

## References

Aguzzi A, Barres BA, Bennett ML (2013) Microglia: scapegoat, saboteur, or something else? Science 339: 156–161

Arber C, Lovejoy C, Wray S (2017) Stem cell models of Alzheimer’s disease: progress and challenges. Alzheimers Res Ther 9: 42

Bateman RJ, Xiong C, Benzinger TL, Fagan AM, Goate A, Fox NC, Marcus DS, Cairns NJ, Xie X, Blazey TM et al (2012) Clinical and biomarker changes in dominantly inherited Alzheimer’s disease. N Engl J Med 367: 795–804

Bellenguez C, Charbonnier C, Grenier-Boley B, Quenez O, Le Guennec K, Nicolas G, Chauhan G, Wallon D, Rousseau S, Richard AC et al (2017) Contribution to Alzheimer’s disease risk of rare variants in TREM2, SORL1, and ABCA7 in 1779 cases and 1273 controls. Neurobiol Aging 59: 220.e221-220.e229

Bennett ML, Bennett FC, Liddelow SA, Ajami B, Zamanian JL, Fernhoff NB, Mulinyawe SB, Bohlen CJ, Adil A, Tucker A et al (2016) New tools for studying microglia in the mouse and human CNS. Proc Natl Acad Sci U S A 113: E1738–1746

Boutej H, Rahimian R, Thammisetty SS, Béland LC, Lalancette-Hébert M, Kriz J (2017) Diverging mRNA and Protein Networks in Activated Microglia Reveal SRSF3 Suppresses Translation of Highly Upregulated Innate Immune Transcripts. Cell Rep 21: 3220–3233

Butovsky O, Jedrychowski MP, Moore CS, Cialic R, Lanser AJ, Gabriely G, Koeglsperger T, Dake B, Wu PM, Doykan CE et al (2014) Identification of a unique TGF-β-dependent molecular and functional signature in microglia. Nat Neurosci 17: 131–143

Carrasquillo MM, Allen M, Burgess JD, Wang X, Strickland SL, Aryal S, Siuda J, Kachadoorian ML, Medway C, Younkin CS et al (2017) A candidate regulatory variant at the TREM gene cluster associates with decreased Alzheimer’s disease risk and increased TREML1 and TREM2 brain gene expression. Alzheimers Dement 13: 663–673

Chakrabarty P, Ceballos-Diaz C, Beccard A, Janus C, Dickson D, Golde TE, Das P (2010) IFN-gamma promotes complement expression and attenuates amyloid plaque deposition in amyloid beta precursor protein transgenic mice. J Immunol 184: 5333–5343

Chakrabarty P, Herring A, Ceballos-Diaz C, Das P, Golde TE (2011) Hippocampal expression of murine TNFα results in attenuation of amyloid deposition in vivo. Mol Neurodegener 6: 16

Chakrabarty P, Li A, Ceballos-Diaz C, Eddy JA, Funk CC, Moore B, DiNunno N, Rosario AM, Cruz PE, Verbeeck C et al (2015) IL-10 alters immunoproteostasis in APP mice, increasing plaque burden and worsening cognitive behavior. Neuron 85: 519–533

Chakrabarty P, Li A, Ladd TB, Strickland MR, Koller EJ, Burgess JD, Funk CC, Cruz PE, Allen M, Yaroshenko M et al (2018) TLR5 decoy receptor as a novel anti-amyloid therapeutic for Alzheimer’s disease. J Exp Med 215: 2247–2264

Chen WT, Lu A, Craessaerts K, Pavie B, Sala Frigerio C, Corthout N, Qian X, Laláková J, Kühnemund M, Voytyuk I et al (2020) Spatial Transcriptomics and In Situ Sequencing to Study Alzheimer’s Disease. Cell 182: 976–991.e919

Chishti MA, Yang DS, Janus C, Phinney AL, Horne P, Pearson J, Strome R, Zuker N, Loukides J, French J et al (2001) Early-onset amyloid deposition and cognitive deficits in transgenic mice expressing a double mutant form of amyloid precursor protein 695. J Biol Chem 276: 21562–21570

Cline EN, Bicca MA, Viola KL, Klein WL (2018) The Amyloid-β Oligomer Hypothesis: Beginning of the Third Decade. J Alzheimers Dis 64: S567–S610

Colon-Perez LM, Ibanez KR, Suarez M, Torroella K, Acuna K, Ofori E, Levites Y, Vaillancourt DE, Golde TE, Chakrabarty P et al (2019) Neurite orientation dispersion and density imaging reveals white matter and hippocampal microstructure changes produced by Interleukin-6 in the TgCRND8 mouse model of amyloidosis. Neuroimage 202: 116138

Conway OJ, Carrasquillo MM, Wang X, Bredenberg JM, Reddy JS, Strickland SL, Younkin CS, Burgess JD, Allen M, Lincoln SJ et al (2018) ABI3 and PLCG2 missense variants as risk factors for neurodegenerative diseases in Caucasians and African Americans. Mol Neurodegener 13: 53

Croft CL, Futch HS, Moore BD, Golde TE (2019) Organotypic brain slice cultures to model neurodegenerative proteinopathies. Mol Neurodegener 14: 45

Cummings J (2019) The National Institute on Aging-Alzheimer’s Association Framework on Alzheimer’s disease: Application to clinical trials. Alzheimers Dement 15: 172–178

Deshpande A, Mina E, Glabe C, Busciglio J (2006) Different conformations of amyloid beta induce neurotoxicity by distinct mechanisms in human cortical neurons. J Neurosci 26: 6011–6018

Dewapriya P, Himaya SW, Li YX, Kim SK (2013) Tyrosol exerts a protective effect against dopaminergic neuronal cell death in in vitro model of Parkinson’s disease. Food Chem 141: 1147–1157

Dickson DW (1997) The pathogenesis of senile plaques. J Neuropathol Exp Neurol 56: 321–339

Dickson DW, Farlo J, Davies P, Crystal H, Fuld P, Yen SH (1988) Alzheimer’s disease. A double-labeling immunohistochemical study of senile plaques. Am J Pathol 132: 86–101

Dobin A, Davis CA, Schlesinger F, Drenkow J, Zaleski C, Jha S, Batut P, Chaisson M, Gingeras TR (2013) STAR: ultrafast universal RNA-seq aligner. Bioinformatics 29: 15–21

Friedman BA, Srinivasan K, Ayalon G, Meilandt WJ, Lin H, Huntley MA, Cao Y, Lee SH, Haddick PCG, Ngu H et al (2018) Diverse Brain Myeloid Expression Profiles Reveal Distinct Microglial Activation States and Aspects of Alzheimer’s Disease Not Evident in Mouse Models. Cell Rep 22: 832–847

Guerreiro R, Wojtas A, Bras J, Carrasquillo M, Rogaeva E, Majounie E, Cruchaga C, Sassi C, Kauwe JS, Younkin S et al (2013) TREM2 variants in Alzheimer’s disease. N Engl J Med 368: 117–127

Haass C, Selkoe DJ (2007) Soluble protein oligomers in neurodegeneration: lessons from the Alzheimer’s amyloid beta-peptide. Nat Rev Mol Cell Biol 8: 101–112

Hammond TR, Dufort C, Dissing-Olesen L, Giera S, Young A, Wysoker A, Walker AJ, Gergits F, Segel M, Nemesh J et al (2019) Single-Cell RNA Sequencing of Microglia throughout the Mouse Lifespan and in the Injured Brain Reveals Complex Cell-State Changes. Immunity 50: 253–271.e256

Hanseeuw BJ, Betensky RA, Jacobs HIL, Schultz AP, Sepulcre J, Becker JA, Cosio DMO, Farrell M, Quiroz YT, Mormino EC et al (2019) Association of Amyloid and Tau With Cognition in Preclinical Alzheimer Disease: A Longitudinal Study. JAMA Neurol

Harold D, Abraham R, Hollingworth P, Sims R, Gerrish A, Hamshere ML, Pahwa JS, Moskvina V, Dowzell K, Williams A et al (2009) Genome-wide association study identifies variants at CLU and PICALM associated with Alzheimer’s disease. Nat Genet 41: 1088–1093

He Y, Yao X, Taylor N, Bai Y, Lovenberg T, Bhattacharya A (2018) RNA sequencing analysis reveals quiescent microglia isolation methods from postnatal mouse brains and limitations of BV2 cells. J Neuroinflammation 15: 153

He Y, Zheng MM, Ma Y, Han XJ, Ma XQ, Qu CQ, Du YF (2012) Soluble oligomers and fibrillar species of amyloid β-peptide differentially affect cognitive functions and hippocampal inflammatory response. Biochem Biophys Res Commun 429: 125–130

Jack CR, Bennett DA, Blennow K, Carrillo MC, Dunn B, Haeberlein SB, Holtzman DM, Jagust W, Jessen F, Karlawish J et al (2018) NIA-AA Research Framework: Toward a biological definition of Alzheimer’s disease. Alzheimers Dement 14: 535–562

Jan A, Adolfsson O, Allaman I, Buccarello AL, Magistretti PJ, Pfeifer A, Muhs A, Lashuel HA (2011) Abeta42 neurotoxicity is mediated by ongoing nucleated polymerization process rather than by discrete Abeta42 species. J Biol Chem 286: 8585–8596

Jan A, Gokce O, Luthi-Carter R, Lashuel HA (2008) The ratio of monomeric to aggregated forms of Abeta40 and Abeta42 is an important determinant of amyloid-beta aggregation, fibrillogenesis, and toxicity. J Biol Chem 283: 28176–28189

Jarosz-Griffiths HH, Noble E, Rushworth JV, Hooper NM (2016) Amyloid-β Receptors: The Good, the Bad, and the Prion Protein. J Biol Chem 291: 3174–3183

Jimenez S, Baglietto-Vargas D, Caballero C, Moreno-Gonzalez I, Torres M, Sanchez-Varo R, Ruano D, Vizuete M, Gutierrez A, Vitorica J (2008) Inflammatory response in the hippocampus of PS1M146L/APP751SL mouse model of Alzheimer’s disease: age-dependent switch in the microglial phenotype from alternative to classic. J Neurosci 28: 11650–11661

Jin SC, Benitez BA, Karch CM, Cooper B, Skorupa T, Carrell D, Norton JB, Hsu S, Harari O, Cai Y et al (2014) Coding variants in TREM2 increase risk for Alzheimer’s disease. Hum Mol Genet 23: 5838–5846

Jin SC, Carrasquillo MM, Benitez BA, Skorupa T, Carrell D, Patel D, Lincoln S, Krishnan S, Kachadoorian M, Reitz C et al (2015) TREM2 is associated with increased risk for Alzheimer’s disease in African Americans. Mol Neurodegener 10: 19

Jonsson T, Stefansson H, Steinberg S, Jonsdottir I, Jonsson PV, Snaedal J, Bjornsson S, Huttenlocher J, Levey AI, Lah JJ et al (2013) Variant of TREM2 associated with the risk of Alzheimer’s disease. N Engl J Med 368: 107–116

Kassambara A, 2020. ggpubr: ’ggplot2’ Based Publication Ready Plots. https://CRAN.R-project.org/package=ggpubr.

Kayed R, Lasagna-Reeves CA (2013) Molecular mechanisms of amyloid oligomers toxicity. J Alzheimers Dis 33 Suppl 1: S67–78

Keren-Shaul H, Spinrad A, Weiner A, Matcovitch-Natan O, Dvir-Szternfeld R, Ulland TK, David E, Baruch K, Lara-Astaiso D, Toth B et al (2017) A Unique Microglia Type Associated with Restricting Development of Alzheimer’s Disease. Cell 169: 1276–1290.e1217

Krasemann S, Madore C, Cialic R, Baufeld C, Calcagno N, El Fatimy R, Beckers L, O’Loughlin E, Xu Y, Fanek Z et al (2017) The TREM2-APOE Pathway Drives the Transcriptional Phenotype of Dysfunctional Microglia in Neurodegenerative Diseases. Immunity 47: 566–581.e569

Kunkle BW, Grenier-Boley B, Sims R, Bis JC, Damotte V, Naj AC, Boland A, Vronskaya M, van der Lee SJ, Amlie-Wolf A et al (2019) Genetic meta-analysis of diagnosed Alzheimer’s disease identifies new risk loci and implicates Aβ, tau, immunity and lipid processing. Nat Genet 51: 414–430

Lambert JC, Heath S, Even G, Campion D, Sleegers K, Hiltunen M, Combarros O, Zelenika D, Bullido MJ, Tavernier B et al (2009) Genome-wide association study identifies variants at CLU and CR1 associated with Alzheimer’s disease. Nat Genet 41: 1094–1099

Lambert JC, Ibrahim-Verbaas CA, Harold D, Naj AC, Sims R, Bellenguez C, DeStafano AL, Bis JC, Beecham GW, Grenier-Boley B et al (2013) Meta-analysis of 74,046 individuals identifies 11 new susceptibility loci for Alzheimer’s disease. Nat Genet 45: 1452–1458

Langfelder P, Horvath S (2008) WGCNA: an R package for weighted correlation network analysis. BMC Bioinformatics 9: 559

Langfelder P, Horvath S (2012) Fast R Functions for Robust Correlations and Hierarchical Clustering. J Stat Softw 46

Lawrence M, Huber W, Pagès H, Aboyoun P, Carlson M, Gentleman R, Morgan MT, Carey VJ (2013) Software for computing and annotating genomic ranges. PLoS Comput Biol 9: e1003118

Li Q, Cheng Z, Zhou L, Darmanis S, Neff NF, Okamoto J, Gulati G, Bennett ML, Sun LO, Clarke LE et al (2019) Developmental Heterogeneity of Microglia and Brain Myeloid Cells Revealed by Deep Single-Cell RNA Sequencing. Neuron 101: 207–223.e210

Li S, Selkoe DJ (2020) A mechanistic hypothesis for the impairment of synaptic plasticity by soluble Aβ oligomers from Alzheimer’s brain. J Neurochem 154: 583–597

Lin L, Desai R, Wang X, Lo EH, Xing C (2017) Characteristics of primary rat microglia isolated from mixed cultures using two different methods. J Neuroinflammation 14: 101

Liu CC, Kanekiyo T, Xu H, Bu G (2013) Apolipoprotein E and Alzheimer disease: risk, mechanisms and therapy. Nat Rev Neurol 9: 106–118

Love MI, Huber W, Anders S (2014) Moderated estimation of fold change and dispersion for RNA-seq data with DESeq2. Genome Biol 15: 550

Morgan M, Pages H, Obenchain V, Hayden N, 2018. Rsamtools: Binary alignment (BAM), FASTA, variant call (BCF), and tabix file import. http://bioconductororg/packages/release/bioc/html/Rsamtoolshtml R package version 1.32.0 ed.

Olah M, Menon V, Habib N, Taga MF, Ma Y, Yung CJ, Cimpean M, Khairallah A, Coronas-Samano G, Sankowski R et al (2020) Single cell RNA sequencing of human microglia uncovers a subset associated with Alzheimer’s disease. Nat Commun 11: 6129

Perlmutter LS, Scott SA, Barrón E, Chui HC (1992) MHC class II-positive microglia in human brain: association with Alzheimer lesions. J Neurosci Res 33: 549–558

Rammes G, Hasenjäger A, Sroka-Saidi K, Deussing JM, Parsons CG (2011) Therapeutic significance of NR2B-containing NMDA receptors and mGluR5 metabotropic glutamate receptors in mediating the synaptotoxic effects of β-amyloid oligomers on long-term potentiation (LTP) in murine hippocampal slices. Neuropharmacology 60: 982–990

Rosario AM, Cruz PE, Ceballos-Diaz C, Strickland MR, Siemienski Z, Pardo M, Schob KL, Li A, Aslanidi GV, Srivastava A et al (2016) Microglia-specific targeting by novel capsid-modified AAV6 vectors. Mol Ther Methods Clin Dev 3: 16026

Sala Frigerio C, Wolfs L, Fattorelli N, Thrupp N, Voytyuk I, Schmidt I, Mancuso R, Chen WT, Woodbury ME, Srivastava G et al (2019) The Major Risk Factors for Alzheimer’s Disease: Age, Sex, and Genes Modulate the Microglia Response to Aβ Plaques. Cell Rep 27: 1293–1306.e1296

Shen L, 2020. GeneOverlap: Test and visualize gene overlaps . R package version 1.26.0.

Sims R, van der Lee SJ, Naj AC, Bellenguez C, Badarinarayan N, Jakobsdottir J, Kunkle BW, Boland A, Raybould R, Bis JC et al (2017) Rare coding variants in PLCG2, ABI3, and TREM2 implicate microglial-mediated innate immunity in Alzheimer’s disease. Nat Genet 49: 1373–1384

Stine WB, Dahlgren KN, Krafft GA, LaDu MJ (2003) In vitro characterization of conditions for amyloid-beta peptide oligomerization and fibrillogenesis. J Biol Chem 278: 11612–11622

Strickland SL, Morel H, Prusinski C, Allen M, Patel TA, Carrasquillo MM, Conway OJ, Lincoln SJ, Reddy JS, Nguyen T et al (2020) Association of ABI3 and PLCG2 missense variants with disease risk and neuropathology in Lewy body disease and progressive supranuclear palsy. Acta Neuropathol Commun 8: 172

Tham CS, Lin FF, Rao TS, Yu N, Webb M (2003) Microglial activation state and lysophospholipid acid receptor expression. Int J Dev Neurosci 21: 431–443

Tseng BP, Esler WP, Clish CB, Stimson ER, Ghilardi JR, Vinters HV, Mantyh PW, Lee JP, Maggio JE (1999) Deposition of monomeric, not oligomeric, Abeta mediates growth of Alzheimer’s disease amyloid plaques in human brain preparations. Biochemistry 38: 10424–10431

van der Lee SJ, Conway OJ, Jansen I, Carrasquillo MM, Kleineidam L, van den Akker E, Hernández I, van Eijk KR, Stringa N, Chen JA, et al (2019) A nonsynonymous mutation in PLCG2 reduces the risk of Alzheimer’s disease, dementia with Lewy bodies and frontotemporal dementia, and increases the likelihood of longevity. Acta Neuropathol 138: 237–250

van Helmond Z, Miners JS, Kehoe PG, Love S (2010) Oligomeric Abeta in Alzheimer’s disease: relationship to plaque and tangle pathology, APOE genotype and cerebral amyloid angiopathy. Brain Pathol 20: 468–480

Vickers JC, Mitew S, Woodhouse A, Fernandez-Martos CM, Kirkcaldie MT, Canty AJ, McCormack GH, King AE (2016) Defining the earliest pathological changes of Alzheimer’s disease. Curr Alzheimer Res 13: 281–287

Villemagne VL, Burnham S, Bourgeat P, Brown B, Ellis KA, Salvado O, Szoeke C, Macaulay SL, Martins R, Maruff P et al (2013) Amyloid β deposition, neurodegeneration, and cognitive decline in sporadic Alzheimer’s disease: a prospective cohort study. Lancet Neurol 12: 357–367

Wang ZX, Tan L, Liu J, Yu JT (2016) The Essential Role of Soluble Aβ Oligomers in Alzheimer’s Disease. Mol Neurobiol 53: 1905–1924

Waters J (2010) The concentration of soluble extracellular amyloid-β protein in acute brain slices from CRND8 mice. PLoS One 5: e15709

Webers A, Heneka MT, Gleeson PA (2020) The role of innate immune responses and neuroinflammation in amyloid accumulation and progression of Alzheimer’s disease. Immunol Cell Biol 98: 28–41

Wickham H (2016) ggplot2: Elegant Graphics for Data Analysis. https://ggplot2.tidyverse.org. Springer-Verlag New York

Young MD, Wakefield MJ, Smyth GK, Oshlack A (2010) Gene ontology analysis for RNA-seq: accounting for selection bias. Genome Biol 11: R14

Zhang Y, Chen K, Sloan SA, Bennett ML, Scholze AR, O’Keeffe S, Phatnani HP, Guarnieri P, Caneda C, Ruderisch N et al (2014) An RNA-sequencing transcriptome and splicing database of glia, neurons, and vascular cells of the cerebral cortex. J Neurosci 34: 11929–11947

